# Discovery of disease-associated cellular states using ResidPCA in single-cell RNA and ATAC sequencing data

**DOI:** 10.1101/2024.12.29.630536

**Authors:** Shaye Carver, Kodi Taraszka, Stefan Groha, Alexander Gusev

## Abstract

To enhance understanding of cellular heterogeneity and disease from single-cell sequencing data, we introduce ResidPCA, a robust method for cell state identification that models cell type heterogeneity. Simulations demonstrate ResidPCA’s efficacy, particularly in complex scenarios, with its accuracy more than four times higher than conventional Principal Component Analysis (PCA) and over three times higher than Non-negative Matrix Factorization (NMF)-based methods in identifying states expressed across multiple cell types. In scRNA-seq data from light-stimulated mouse visual cortex cells, ResidPCA captures stimulus-driven variability with an accuracy more than five times higher than NMF methods. In single nucleus data from an Alzheimer’s disease cohort, ResidPCA identified 44 snATAC-based and 42 snRNA-based states. 30 snATAC states were significantly enriched for Alzheimer’s disease heritability and were often more significantly enriched than established cell types such as microglia. The ResidPCA-based snATAC state most significantly enriched for Alzheimer’s disease heritability further elucidates a recently identified mechanism involving the neuron-ODC-microglial axis. This state links early amyloid production in neurons and oligodendrocytes with later-stage microglial activation and immune response, driving Alzheimer’s disease progression. These results demonstrate ResidPCA’s ability to reveal additional biological variation in single-cell data and uncover disease-relevant cell states.

## Introduction

Single-cell omic measurements have been used to dissect the heterogeneity of cell populations at unprecedented resolution, with a focus on cell type inference^1,2^. The integration of genome-wide association studies with -omic data has shown that many diseases, including neurodegenerative disorders, are linked to and even modulate disease through a diverse range of cell types^3–8^. Despite their disease relevance, only a small fraction of heritability is mediated by existing gene expression measurements due to the complexity of regulatory effects across tissues and cell types^6^. Recent studies have attempted to implicate the missing heritability by moving beyond canonical cell types to more granular cell “states”, which capture the underlying functional mechanisms of tissues and cell types^5,9–14^. These causal mechanisms provide deeper insights into the biological changes driving disease phenotypes and aid in identifying more precise druggable targets compared to a broad cell type approach^14^.

Previous efforts in identifying cell states have largely relied on methodologies borrowed from cell type inference. These approaches can lack sensitivity to both identify all active states within a dataset and to connect them to specific traits or diseases. For instance, many studies adopt unsupervised clustering methods to identify cell states, initially designed for discrete cell type identification^7,15,16^. This strategy may erroneously partition continuous cell states into discrete clusters, restricting each cell to a singular state—a biologically plausible approach for identifying cell types but not states, where cells are capable of occupying multiple states concurrently^11^. This approach likewise restricts the exploration of cellular states to those within a specific cell type, thus limiting the discovery of potential shared states that span multiple cell types.

To address these limitations, methods have been introduced for inferring continuous cell representations using various dimensionality reduction implementations. Such methods include Non-negative Matrix Factorization (NMF)^17^, Consensus Non-negative Matrix Factorization (cNMF)^12^, Principal Component Analysis (PCA)^14,18^, Uniform Manifold Approximation and Projection (UMAP) with unsupervised clustering^15,16,19–21^, and co-expression analyses^22^. However, no existing methods model cell type heterogeneity–the effect of cell type on expression–to accurately estimate cell states that are independent of cell type. Additionally, the sensitivity of existing methods can be severely limited by cell type heterogeneity, which is the largest source of biological variance in single cell datasets^23,24^. While some techniques attempt to address cell type-driven noise by applying PCA to individual cell types (which we refer to as Iterative PCA), this compromises statistical power by reducing sample size and may miss states that are rare or present in multiple cell types^14^.

Here, we develop and benchmark our method, Residual PCA (ResidPCA). This method addresses cell type heterogeneity by utilizing known cell type labels to efficiently denoise single-cell datasets, resulting in the elimination of irrelevant and noisy biological signals that can obscure accurate cell state identification. Through analyses of simulated and real data, we demonstrate that ResidPCA exhibits higher sensitivity than existing approaches, particularly in scenarios featuring many cell types, states spanning multiple cell types, or rare cell states. In an application of ResidPCA to human brain derived single-nuclei RNA (snRNA-seq) and ATAC-seq (snATAC-seq) data, we identify 86 total states accounting for a higher fraction of Alzheimer’s disease (AD) heritability compared to previously established cell type enrichments and map states to their respective biologically relevant pathways. We introduce an efficient implementation of ResidPCA, 3.5 times faster and 5 times more resource-efficient than current state-of-the-art methods.

## Results

### ResidPCA overview

Here, we propose Residual Principal Component Analysis (ResidPCA), a method to identify a denoised set of latent cell states from single cell data, that is efficient and robust to cell type heterogeneity. The procedure for ResidPCA is as follows: it identifies cell type from single-cell expression data by leveraging established approaches (e.g. marker genes or projection from an atlas), it then regresses out cell type-driven expression, and finally it applies PCA to the residualized matrix corresponding to state driven expression. This method effectively denoises the data by removing cell type-driven expression, which can obscure accurate cell state estimation due to variability or noise in gene expression associated with cell identity. As a result, ResidPCA can identify a diverse range of cell states—both rare and common—that are independent of cell type and may be unique to a single cell type or span multiple cell types.

The generative model underlying ResidPCA assumes the log-normalized TP10k (Transcripts Per 10,000) gene expression data can be represented as a mixture of two multivariate Gaussian distributions with identical variances: one reflecting state-driven expression and the other capturing cell type-driven expression. Specifically, the model assumes:

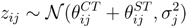

where *z*_*ij*_ represents the normalized gene expression for gene *j* in cell 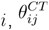, denotes the cell type-driven expression component, 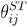 denotes the state-driven expression component, and 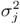 denotes the variance for gene *j*.

To estimate the cell type-driven expression component for gene *j* in cell 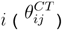,ResidPCA decomposes it as the inner product of the cell type indicator for cell 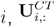, (the *i*-th row), and the normalized mean expression vector for gene *j* in each cell type, 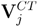 (the *j*-th column) , which indicates how active a gene is in each cell type expression profile:

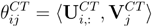

The state-driven component for gene *j* in cell 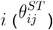 is estimated similarly, using the inner product of the cell state embeddings for cell 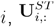 (the *i*-th row), which represent the state occupancy of each cell and the cell state gene loadings for gene 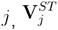 which indicate how active a gene is in each state:

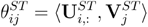

Thus, the normalized expression data *z*_*ij*_ can be reparameterized as:

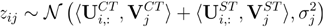

First, ResidPCA estimates the cell-type driven expression matrix, 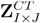 (where *I* is the total number of cells and *J* is the total number of genes), and the state-driven expression matrix,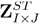 from the normalized and standardized TP1Ok gene expression matrix Z_*I* ×*J*_. For each gene *j*, the cell type-specific expression component 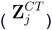 is estimated via linear regression. This regression models the relationship between the cell type indicator matrix, U^*CT*^, and the total normalized and standardized TP1Ok expression for gene *j*(Z_*j*_ ). The regression coefficients,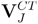 , represent the cell-type driven mean expression of gene *j* across all cell types:

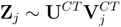

To compute the state-driven expression matrix, we subtract the estimated cell type-driven expression component—derived as the reconstruction estimate from the above regression—from the total TP1Ok-normalized expression. This yields an expression component that is uncorrelated with cell type and represents the additive state-driven expression component of gene *j* across all states:

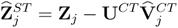

Where 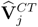 represents the estimated regression coefficients or betas outputted by the regression for a given gene *j*.

In the final step of ResidPCA, the estimated matrix representing the total state-driven expression across all states 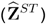 is decomposed into individual state-specific components. PCA is applied to the covariance matrix of 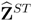 to obtain the state-driven cell embeddings, 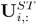, for each cell *i*, and the state-driven gene loadings,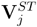 , for each gene j, along with the associated eigenvalues ordering states by total variance explained. ResidPCA generates a set of states that captures both how strongly each cell expresses each state ( U^*ST*^) and which genes regulate each state (V^*ST*^ ).

### Benchmarking ResidPCA in simulation

We compare ResidPCA to multiple well-established dimensionality reduction methods/approaches: Standard PCA, which identifies primary and global sources of variation in data by projecting it onto new, orthogonal axes representing the most significant directions of variance^25^, Non-negative Matrix Factorization, NMF, which produces non-orthogonal components that are non-negative and represent additive combinations of the original single cell count data^26^, Consensus Non-negative Matrix Factorization (cNMF), an extension of NMF, which improves the stability of NMF by consolidating the latent representations across multiple runs of NMF^12^, and Scaled NMF, which regresses out covariates from a positive count matrix and enforces non-negativity by setting any negative values to zero before inputting the matrix into the NMF algorithm^27,28^. And finally, Uniform Manifold Approximation and Projection (UMAP) is also used, which is the traditional method used for visualizing single cell data that aims to preserve both global and local structure in the data^29^. Although recent findings indicate that UMAP does not consistently preserve these high-dimensional structures in its two dimensional representation, we include it as a measure of what users might expect to see from typical visual inspection of the data^30^.

We systematically evaluated the accuracy of ResidPCA compared to well-established methods in recovering cell states from simulated data. Single-cell datasets were generated using realistic parameters learned from real data (see **Fig. 1 A, Methods** and **Supplementary Fig. 1**) and simulated to reflect two scenarios: a state expressed in a subset of the total cell population across all cell types (**Fig. 1 B**) and a state exclusively expressed in a subset of a single cell type (**Fig. 1 C**). Simulations involved a small set of total cell types (7) or a larger set (100) to assess method performance under varying levels of cell type heterogeneity. In **Fig. 1 B**, we illustrate an example of a simulated state that is expressed across all seven cell types. Notably, ResidPCA demonstrates superior sensitivity by detecting this state in its first PC (Principal Component), while Standard PCA identifies the state in a later PC (PC7) (**Fig. 1 D**). Likewise, in **Fig. 1 C**, we present an example of a simulated state that spans only one of the seven cell types. Here, we again see ResidPCA has more sensitivity by identifying the state in the second PC, whereas Standard PCA fails to capture it in any single PC (**Fig. 1 E**).

**Fig 1:**
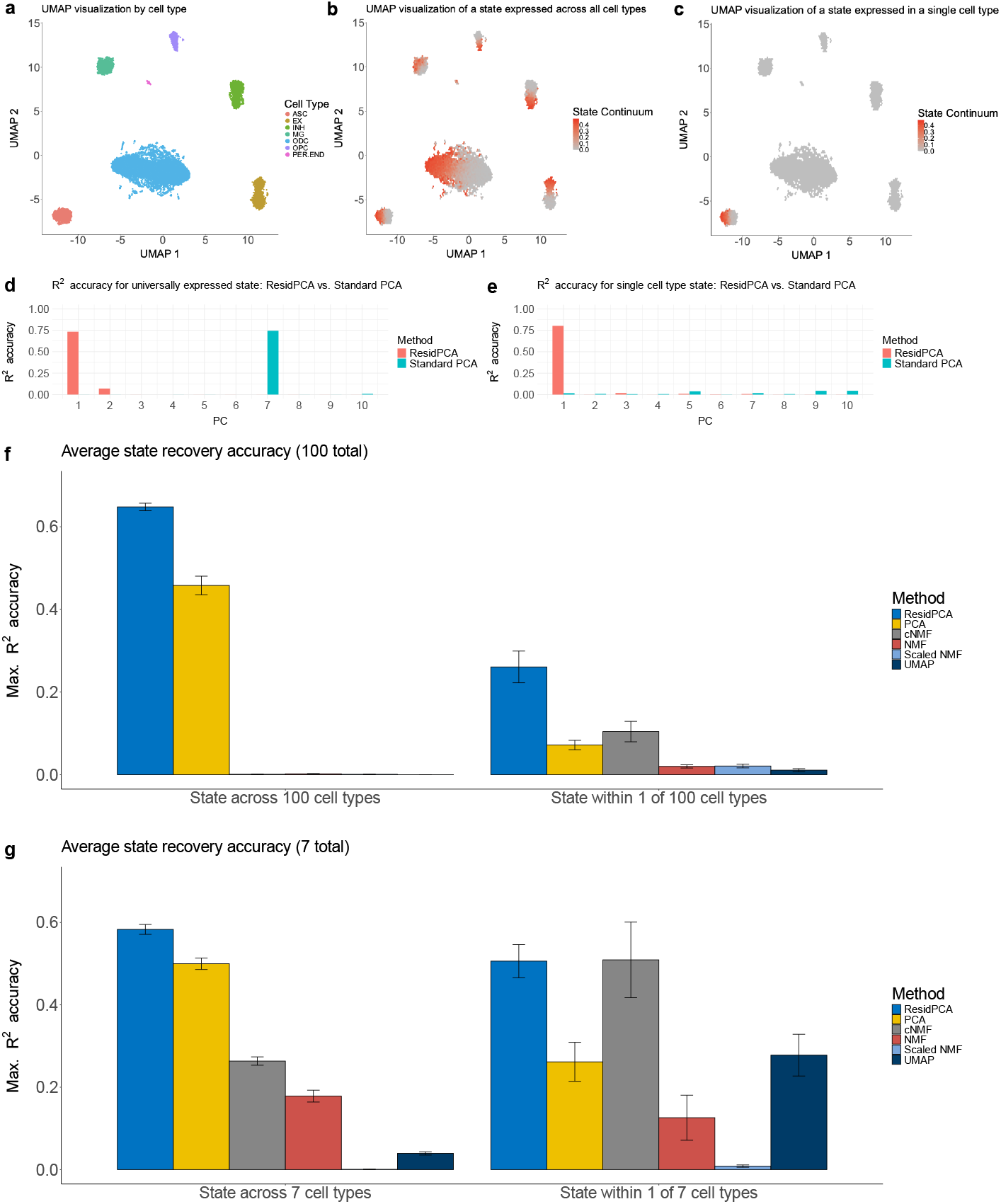
UMAP visualizations of example simulations, colored by (**a**) cell type, (**b**) a state expressed across all seven cell types, and (**c**) a state expressed exclusively in 1/7 cell types. Count matrices were simulated by varying the fraction of genes and cells expressing the state across a grid of parameters. R^2^ metrics were averaged for each parameter set across ten seeds until the standard error of the R^2^ metrics fell below 0.085. (**d, e**) Bar plots showing R^2^ accuracy between the ground truth simulated state continuum and the corresponding PC embedding identified by ResidPCA and Standard PCA, for (**d**) a state expressed across all seven cell types and (**e**) a state expressed within a single cell type. Note: to ensure accurate comparisons, R^2^ correlations were calculated after subsetting both the ground truth state and the recovered embedding to include only the cell type(s) capable of expressing the state. This approach prevents inflated accuracy that could arise from correlations between cell types and the given state. (**f, g**) Bar plots of the average accuracy of each method in capturing the true simulated state continuum, averaged across different parameter settings. Standard errors are plotted across parameter settings. These settings vary in the number of genes and cells involved in the state. Each parameter configuration was replicated 10 times. Averaged maximum R^2^ accuracy in state recovery is shown for states expressed across all cell types (left) and within a single cell type (right) for simulations with (**f**) 100 and (**g**) seven total cell types.

For each method, we then quantified the accuracy of the state detection in each simulation by computing the maximum R^2^ between each inferred component and the true state continuum. The maximum was used (as opposed to the R^2^ with the “first” component) because some methods produce an arbitrary ordering of inferred components. For each simulated state, only cell types in which the state was putatively active were considered when quantifying accuracy, to distinguish cell state from cell type detection, meaning that if the state was expressed in a single cell type, both the embeddings and the ground truth state continuum were subsetted to that specific cell type, and correlations were computed accordingly. Conversely, if the state was expressed across all cell types, the unsubsetted embeddings and state continuum were used to compute the correlations.

ResidPCA consistently demonstrated higher accuracy compared to other methods in simulations, particularly when cell type heterogeneity was high or states spanned multiple cell types. We simulated an intentionally high cell type heterogeneity scenario consisting of 100 total cell types; ResidPCA had higher accuracy than all other methods, regardless of whether states were expressed within a single cell type or across all cell types (**Fig. 1 F**). For simulations with states active across all cell types, ResidPCA achieved an average R^2^ of 0.65 (CI: 0.63-0.67), compared to an average R^2^ of 0.46 (CI: 0.42-0.50) for Standard PCA and R^2^ < 0.0021 (upper CI < 0.0029) for all other methods (cNMF, NMF, UMAP) (**Fig. 1 F**). For simulations with states active in 1/100 total cell types, all methods had reduced performance compared to ResidPCA, with a mean R^2^ of 0.26 (CI: 0.19-0.34) compared to a mean R^2^ of 0.10 (CI: 0.06-0.15) for cNMF and 0.072 (CI: 0.05-0.09) for Standard PCA (**Fig. 1 F**). Across all methods, overall performance improved with increased state abundance (defined as states with a high fraction of cells and genes expressing the state), exemplified by the average difference in accuracy between an abundant versus rare cell state (ResidPCA ΔR^2^ = 0.63, Standard PCA ΔR^2^ = 0.70, cNMF ΔR^2^ = 7.8×10^-4^, NMF ΔR^2^ = 6.4×10^-3^, scenario: one state expressed across 100 cell types) (**Supplementary Fig. 1**). ResidPCA was consistently more sensitive compared to alternative methods, requiring fewer cells and genes expressing the simulated state to detect states effectively; for instance, in simulations where the state was present within 1/100 cell types, and with a fixed total number of genes expressing the state, Standard PCA (the next best performing method) required over 4.7 times more cells to attain a similar accuracy as ResidPCA (**Supplementary Fig. 1**).

In simulations with lower cell type heterogeneity or a smaller number of cell types (7), ResidPCA was still the most accurate method, although differences in performance between ResidPCA and Standard PCA were less pronounced (**Supplementary Fig. 1**). In simulations with a state spanning all seven cell types, ResidPCA performed better on average than Standard PCA (mean R^2^ of 0.58 (CI: 0.56-0.61) and 0.50 (CI: 047-0.53), respectively) and both methods substantially outperformed cNMF (mean R^2^ of 0.26 (CI: 0.24-0.28)) (**Fig. 1 G**). ResidPCA outperformed Standard PCA when the percentage of genes in the state was low or the state was rarer, but was otherwise comparable (**Supplementary Fig. 1**). In simulations with a state spanning 1/7 cell types, ResidPCA had comparable performance to cNMF (mean R^2^ of 0.51 (CI: 0.43-0.59) and 0.51 (CI: 0.33-0.69), respectively) and substantially outperformed Standard PCA (mean R^2^ of 0.26 (CI: 0.17-0.35)). Overall performance again improved with increased state abundance in states across seven cell types, exemplified by the average difference in accuracy between an abundant versus rare cell state per method (ResidPCA ΔR^2^ = 0.83, Standard PCA ΔR^2^ = 0.76, cNMF ΔR^2^ = -0.084, NMF ΔR^2^ = 0.24, scenario: one state expressed across seven cell types) (**Supplementary Fig. 1**).

We next evaluated the ability of various methods to capture cell state signals by examining two metrics: the joint adjusted R^2^ across all significant recovered components, which indicates whether the cell state signal is detected at all, and the maximum R^2^ (described previously), which assesses whether the method accurately consolidates the individual cell state signal into a single component rather than dispersing it across multiple components. For abundant states—defined as those with a high proportion of cells and genes expressing the state—all methods demonstrated high joint adjusted R^2^, indicating their ability to accurately identify these states across all learned components. Among them, cNMF slightly outperformed ResidPCA and Standard PCA, in a state spanning seven cell types (cNMF joint adj. R^2^ = 0.68, ResidPCA joint adj. R^2^ = 0.64, PCA joint adj. R^2^ = 0.52) (**Supplementary Fig. 1**). However, in capturing the true state with a single component, NMF-based methods exhibited decreased accuracy compared to PCA-based methods (ResidPCA max. R^2^ = 0.58, PCA max. R^2^ = 0.50, cNMF max. R^2^ = 0.26) (**Supplementary Fig. 1**). This suggests that NMF-based methods are more prone to splitting single states into multiple components than PCA-based methods. Additionally, the two UMAP dimensions generally do not capture a linear correlation with cell state, except in straightforward cases where the state is abundant, cell type heterogeneity is low, and the state is confined to a single cell type (**Supplementary Fig. 1**).

Overall, ResidPCA most accurately recovered simulated states in the majority of instances compared to other methods, particularly in single-cell datasets with a high number of cell types or when states spanned all cell types. Notably, there were no scenarios where ResidPCA was significantly less accurate than Standard PCA, indicating that the former can serve as a direct substitute for the latter when cell types are known. There were a small number of scenarios when ResidPCA was less accurate than cNMF, primarily when the number of cells and the number of genes in a state was small and all methods had low accuracy.

### Benchmarking ResidPCA against state-of-the-art methods in data with experimentally induced states

In order to assess the effectiveness of methods in real data, we turned to a scRNA-seq dataset with an experimentally induced state. In a study from Hvratin et al., 2018, (previously used to benchmark cNMF^12^), mice were exposed to varying durations of light, shown to lead to a transcriptional state change in the mouse visual cortex^31^. We used the amount of light exposure (hours) as a proxy for the true cell state when evaluating performance of the six state detection methods evaluated in simulations (ResidPCA, Standard PCA, cNMF, NMF, Scaled NMF, and UMAP). Light exposure may also influence the relative cell type abundances; therefore, to quantify the ability of the inferred components to capture light-induced states rather than cell type variation, we calculated R^2^ accuracy with light exposure within each classified cell type. We additionally evaluated the impact of using a small subset of (3000) variable genes versus nearly all (20,000) genes on performance, which could not be tested in simulations without imposing specific biological assumptions.

ResidPCA consistently outperformed other methods in recovering state changes across cell types, particularly vascular cell types (endothelial & smooth muscle cells [grouped cell types] and mural cells), which were found to be among the most transcriptionally affected by light exposure, second only to excitatory neurons in the original publication^31^. Using 3,000 variable genes, ResidPCA achieved a max R^2^ of 0.24 (CI: 0.23-0.24) for endothelial & smooth muscle cells, and a max R^2^ of 0.16 (CI: 0.15-0.17) for mural cells; whereas cNMF and NMF showed negligible detection in both cell types (**Fig. 2 A**). The performance of ResidPCA and cNMF were more similar for excitatory neurons, with max R^2^ of 0.044 and 0.046 respectively (with negligible confidence intervals) (**Fig. 2 A**). However, ResidPCA achieved a substantially higher joint adjusted R^2^ of 0.17 compared to 0.077 for cNMF. This supports the hypothesis that excitatory neurons, the cell type most affected by light, induce multiple state changes in response to light exposure that can be captured across multiple distinct components. Our results differ from prior analyses of this data, which only assessed performance after subsetting the input data to excitatory neurons and interneurons (whereas we use the full dataset, including all cell types as input)^12^. Consequently, cNMF may be less effective for initial exploration of cell states when the target cell type(s) expressing a state are unknown, whereas ResidPCA excels at detecting these novel states. In all analyses, ResidPCA outperformed Standard PCA and cNMF outperformed NMF (**Fig. 2 A**). Our findings indicate that ResidPCA effectively identifies unique light-induced states activated in each cell type and can properly recover distinct cell-type specific states into separate principal components (**Supplementary Note 1 & 2**). This observation aligns with the conclusions of Hrvatin et al.^31^ that each cell type activates a distinct cell state or set of genes in response to light exposure.

**Fig 2:**
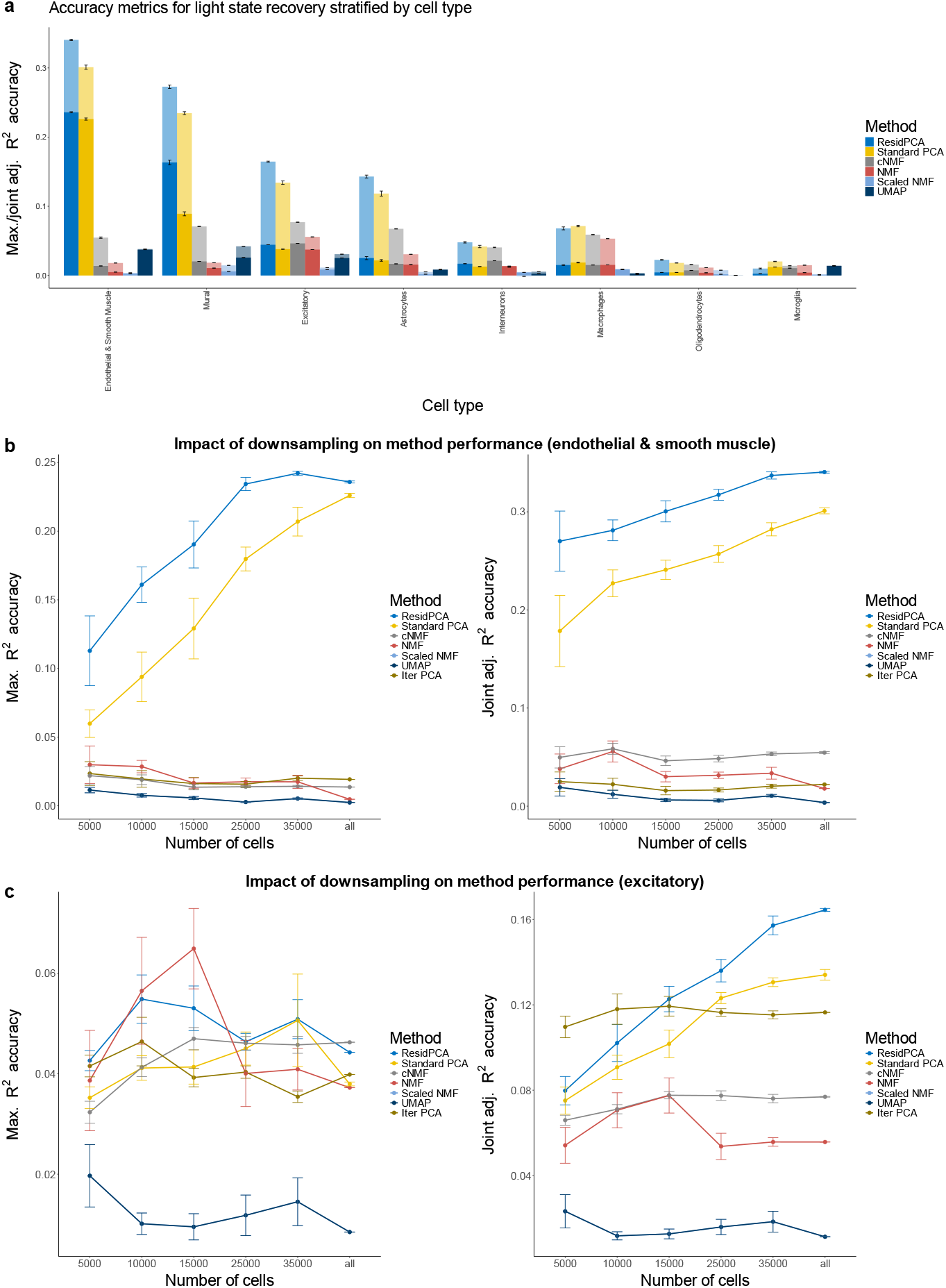
**(a)** Bar plots illustrate two accuracy metrics used in this figure: (1) the maximum R^2^ correlation between all significantly recovered states by a given method and light exposure duration, represented by solid bars, and (2) the joint adjusted R^2^ correlation between all significantly recovered states per method and light exposure duration across each cell type, shown as transparent bars. (**b**,**c**) Accuracy plots were generated for each accuracy metric (left: maximum R^2^, right: joint adj. R^2^) with incremental and random downsampling of total cells (x-axis). Downsampling analysis is shown for endothelial & smooth muscle (**b**) and excitatory neurons (**c**), respectively. Each downsampling had 5 replicates and was averaged across replicates. Standard errors are plotted across replicates. Analyses were performed using 3000 variable genes. Note: the Bayesian Information Criterion (BIC) cutoff^79^ was used to identify the number of significant states in PCA based methods while stability/error maximization^12^ was used to identify the number of significant states in NMF based methods (see Methods).

Next, we downsampled the number of cells to determine how each method performed under more data-limited or underpowered scenarios. ResidPCA was the leading method in 54% of downsampled analyses in terms of max R^2^ and in 81% of downsampled analyses in terms of joint adjusted R^2^. ResidPCA was substantially more sensitive for endothelial & smooth muscle cell types, where Standard PCA required 2.5x as many cells to detect light induced states at the same max R^2^ accuracy while NMF based methods never achieved as high of accuracy as PCA based methods (**Table 2, 1st row**) (**Fig. 2 B**). Likewise for excitatory neurons, Standard PCA required 1.7x as many cells while cNMF never achieved the same R^2^ accuracy as ResidPCA (**Table 2, 2nd row**) (**Fig. 2 C**). Finally, Iterative PCA consistently underperformed in identifying light-induced, cell type-specific states. It never matched the accuracy of ResidPCA, even with an increased number of cells or samples (**Table 2, 4th & 5th row**) (**Fig. 2 B & C**). This underperformance of Iterative PCA likely arises from the fact that using only a subset of the data reduces sensitivity for states that span multiple cell types (even if they are dominant in one cell type). We also investigated the sensitivity of each method to the number of genes included in the analysis. ResidPCA exhibited the smallest variability in accuracy relative to the number of variable genes included in the analysis, but no single optimal gene count emerged as universally the best (**Supplementary Fig. 2, Supplementary Note 3**).

**Table 1:** Overview and comparison of various benchmarked methods used for detecting cell states.

| Method | Description |
| --- | --- |
| ResidPCA | An extension of Principal Component Analysis (PCA) tailored for identifying cell states in single-cell data, where cell type effects are regressed out from the data matrix, allowing the recovered orthogonal components to represent a denoised set of cell states not confounded by cell type driven noise in expression. |
| Principal Component Analysis (PCA) [standard] | A dimensionality reduction method that transforms a matrix into a set of orthogonal components, capturing the maximum variance in the data with fewer dimensions. |
| Iterative PCA | PCA applied iteratively to a subset of the single-cell data specific to each cell type, effectively identifying a set of cell type specific states for each individual cell type. |
| Non-negative Matrix Factorization (NMF) | A factorization method that decomposes a matrix into non-negative components. |
| Consensus Non-Negative Matrix Factorization (cNMF) | An extension of NMF that incorporates multiple factorizations of NMF to identify stable and robust components in data. |
| Scaled NMF | An extension of NMF that enables the regression or removal of covariates by first regressing out the covariate effects from single-cell data and then setting any resulting negative values to zero to maintain non-negativity. |
| Uniform Manifold Approximation and Projection (UMAP) | A dimensionality reduction method that preserves the local structure of the data while reducing its dimensionality for visualization purposes. |

| Feature | ResidPCA | Standard PCA | Iterative PCA | NMF | cNMF | Scaled NMF | UMAP |
| --- | --- | --- | --- | --- | --- | --- | --- |
| Linear vs. Nonlinear | Linear | Linear | Linear | Linear | Linear | Linear | Non-linear |
| Orthogonality Constraint | Yes | Yes | Yes | No | No | No | No |
| Non-Negativity Constraint | No | No | No | Yes | Yes | Yes | No |
| Preservation of Global Structure | High | High | High | Moderate | Moderate | Moderate | Low |
| Scalability to Large Datasets | High | High | High | Moderate | Moderate | Moderate | High |
| Applied to entire dataset | Yes | Yes | No (applied iteratively to each cell type) | Yes | Yes | Yes | Yes |

**Table 2:**
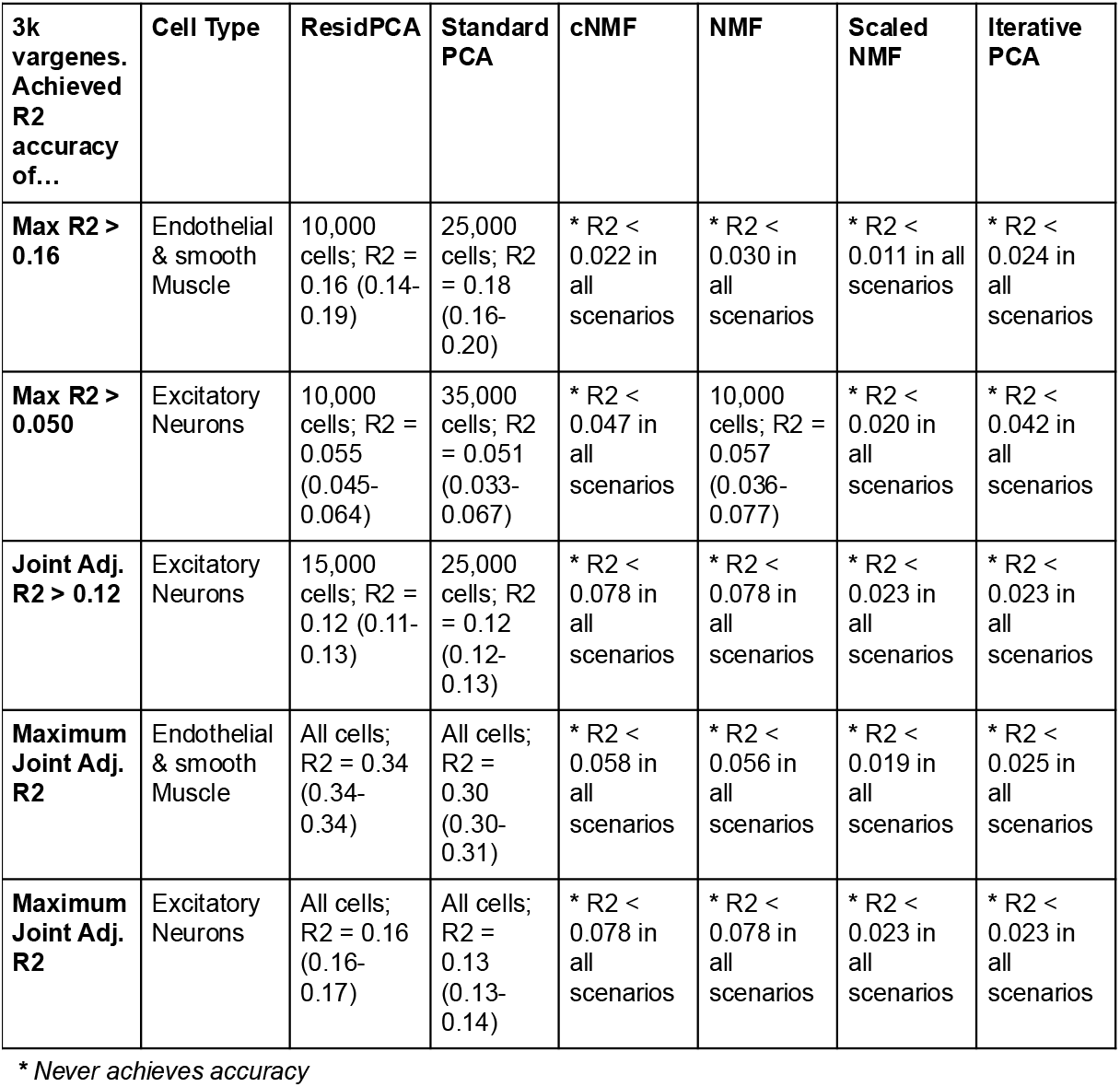
Comparison of the number of cells needed to achieve a specific performance accuracy level across various methods. The first two rows indicate the number of cells required to reach a maximum R^2^ of 0.16 for endothelial & smooth muscle cells, and 0.050 for excitatory neurons, respectively. The third row shows the number of cells necessary to achieve a joint adjusted R^2^ greater than 0.12 for excitatory neurons. The final two rows illustrate the maximum joint adjusted R^2^ attained by each method in endothelial & smooth muscle cells, respectively. An asterisk (*) denotes methods that did not reach the specified accuracy level.

| 3k<br>vargenes.<br>Achieved<br>R2<br>accuracy<br>of... | Cell Type | ResidPCA | Standard<br>PCA | cNMF | NMF | Scaled<br>NMF | Iterative<br>PCA |
| --- | --- | --- | --- | --- | --- | --- | --- |
| Max R2 > 0.16 | Endothelial & smooth Muscle | 10,000 cells; R2 = 0.16 (0.14-0.19) | 25,000 cells; R2 = 0.18 (0.16-0.20) | * R2 < 0.022 in all scenarios | * R2 < 0.030 in all scenarios | * R2 < 0.011 in all scenarios | * R2 < 0.024 in all scenarios |
| Max R2 > 0.050 | Excitatory Neurons | 10,000 cells; R2 = 0.055 (0.045-0.064) | 35,000 cells; R2 = 0.051 (0.033-0.067) | * R2 < 0.047 in all scenarios | 10,000 cells; R2 = 0.057 (0.036-0.077) | * R2 < 0.020 in all scenarios | * R2 < 0.042 in all scenarios |
| Joint Adj. R2 > 0.12 | Excitatory Neurons | 15,000 cells; R2 = 0.12 (0.11-0.13) | 25,000 cells; R2 = 0.12 (0.12-0.13) | * R2 < 0.078 in all scenarios | * R2 < 0.078 in all scenarios | * R2 < 0.023 in all scenarios | * R2 < 0.023 in all scenarios |
| Maximum Joint Adj. R2 | Endothelial & smooth Muscle | All cells; R2 = 0.34 (0.34-0.34) | All cells; R2 = 0.30 (0.30-0.31) | * R2 < 0.058 in all scenarios | * R2 < 0.056 in all scenarios | * R2 < 0.019 in all scenarios | * R2 < 0.025 in all scenarios |
| Maximum Joint Adj. R2 | Excitatory Neurons | All cells; R2 = 0.16 (0.16-0.17) | All cells; R2 = 0.13 (0.13-0.14) | * R2 < 0.078 in all scenarios | * R2 < 0.078 in all scenarios | * R2 < 0.023 in all scenarios | * R2 < 0.023 in all scenarios |

*\* Never achieves accuracy*

### ResidPCA uncovers diverse state types: from broad cell type associations to unique or unassigned states

To demonstrate their utility to discover novel biology, we applied PCA-based methods, including ResidPCA, to a large multi-modal, single-nuclei dataset from late-stage Alzheimer’s Disease (AD) patients, which we re-analyzed and QC’ed (**Fig. 3 A**, see Methods)^15^. We chose this dataset due to several key factors. First, single-cell studies have increasingly focused on brain-derived data because of the brain’s vast cellular heterogeneity and intricate spatial organization^9,15^. Recent advances have identified over 3,000 distinct cell types and states in the human brain, underscoring its complexity^9,20,32,33^. Additionally, the growing prevalence of neuroinflammatory disorders, particularly AD, coupled with the known polygenicity and high heritability of complex brain disorders like AD, as highlighted by large GWAS studies, makes this dataset particularly relevant for understanding the molecular and cellular mechanisms underlying AD^34^.

**Fig 3:**
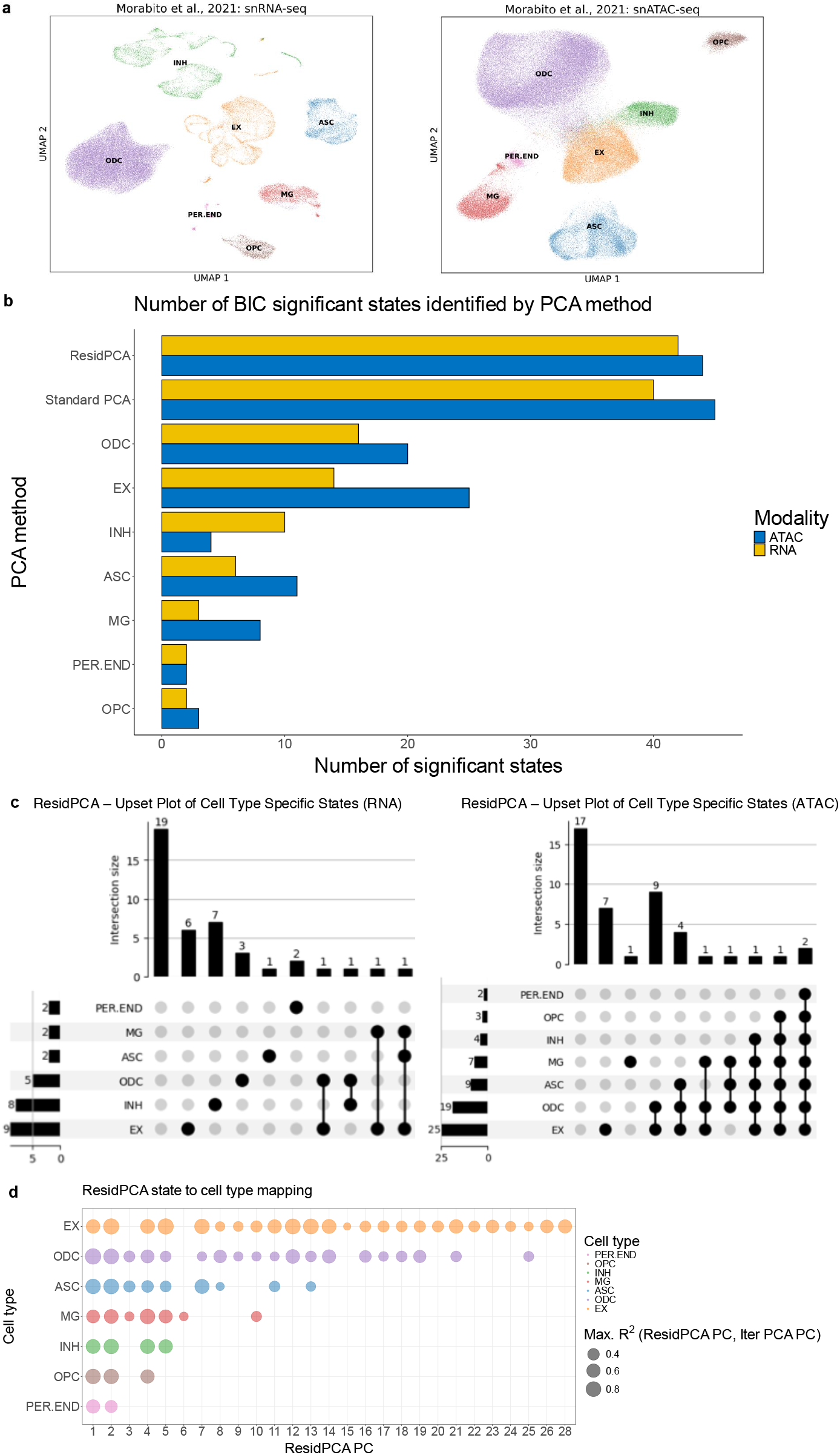
**(a)** UMAP visualizations of re-QC’d data from Morabito et al., 2021^15^, showcasing snRNA-seq gene expression (left) and snATAC-seq peak expression (right). ResidPCA and established PCA methods, including Standard and Iterative PCA, were applied separately to each dataset. **(b)** A bar plot displaying the number of Bayesian Information Criterion (BIC) significant states identified per modality for each PCA method. The y-axis labels containing cell type abbreviations indicate Iterative PCA performed exclusively on data from the corresponding cell type. **(c)** Upset plots illustrating the frequency and distribution of cell types expressing ResidPCA-recovered states across RNA (left) and ATAC (right) data, categorized based on whether the states span one, multiple, all, or no cell types. **(d)** A bubble plot illustrating the maximum R^2^ correlation between each ResidPCA state (x-axis) and all BIC significant Iterative PCA cell states per cell type. Only R^2^ values exceeding 0.20 are plotted. The plot highlights the association of each ResidPCA-based state with specific cell types, as detected by Iterative PCA (see Methods). Bubble sizes correspond to the magnitude of the correlation. Abbreviations: ODC, oligodendrocytes; EX, excitatory neurons; INH, inhibitory neurons; ASC, astrocytes; MG, microglia; PER.END, pericytes and endothelial cells; OPC, oligodendrocyte precursor cells.

When analyzing sc/snATAC-seq data, which is typically very sparse, peak data was mapped to the nearest gene of interest, since this has previously been shown to improve the signal-to-noise ratio in this modality^35^ while enabling consistent state comparisons with sc/snRNA-seq. ResidPCA conditioned out the effect of cell type on gene expression, removing approximately 15 and 20% of the variation in snATAC and snRNA expression, respectively, in the Morabito et al., 2021 datasets (see Methods). The Bayesian Information Criterion (BIC) adapted for PCA (see Methods), was applied to ResidPCA-identified states, revealing 42 and 44 significant cell states in RNA and ATAC-seq data, respectively (**Fig. 3 B, Supplementary Fig. 3**). The BIC-significant states identified by ResidPCA explained 15% of the cumulative variance in the residualized matrix for the RNA dataset and 6% for the ATAC dataset (**Supplementary Fig 3**).The BIC method yielded many more significant states for both RNA and ATAC than the elbow plot method (RNA: 4 states, ATAC: 5 states), suggesting much higher dimensionality in the data than typically considered (**Fig. 3 B, Supplementary Fig. 3**). Among the ResidPCA-identified ATAC-based states, 2 states spanned all cell types (states correlated with all cell types), 17 states exhibited specificity by spanning multiple but not all cell types, 8 states were expressed in only a single cell type, and 17 states were cell type agnostic-newly and uniquely identified by ResidPCA-or rare in one or more cell types, making them challenging to detect using cell type-specific methods like Iterative PCA (**Fig. 3 C**) (see Methods).

Since ATAC data was the only modality that generated cell states expressed in all known cell types, we functionally interpreted those two states, which corresponded to the top PCs identified by ResidPCA, explaining the greatest variance in ATAC expression (**Fig. 3 C, D**). Using Gene Set Enrichment Analysis (GSEA), we found that PC1 was associated with an interconnected set of pathways that contribute to disease pathology through various mechanisms including **GABA receptor complex** (FDR p-value: p < 1.2e-05, Normalized Enrichment Score (NES): 3.03), metabolic pathways like **steroid hormone biosynthesis** (FDR p-value: p < 1.2e-05, NES: 2.6) and **glucuronidation** (FDR p-value: p < 1.2e-05, NES: 2.54), and immune response pathways including **type I interferon receptor binding** (FDR p-value: p < 1.2e-05, NES: 2.7). Dysregulation of GABA receptor complexes can lead to imbalanced neurotransmission and increased excitotoxicity, which may exacerbate neurodegenerative processes in AD^36,37^ while alterations in steroid hormone biosynthesis impact brain function and neuroprotection, potentially interacting with neurotransmitter systems and inflammatory responses^38^. Both neurons and astrocytes produce cholesterol, a hormone strongly implicated in AD, supporting the idea that the state identified by ResidPCA involves multiple cell types^39^. Impaired glucuronidation affects the detoxification and clearance of neurotoxic substances like amyloid-beta, which can exacerbate neuroinflammation and influence hormonal and neurotransmitter pathways^40,41^ while chronic inflammation mediated by type I interferons promotes neuroinflammation mediated by microglia and endothelial cells, which might disrupt both hormonal balance and neurotransmitter systems, creating a feedback loop that accelerates AD progression^42,43^. Thus, these pathways collectively contribute to AD through a variety of cell types, by affecting neurotransmission, neuroinflammation, and the metabolism of neurotoxic substances.

In contrast, PC2 predominantly reflects immune response pathways, including **peptide antigen assembly with major histocompatibility complex (MHC) protein complex** (FDR p-value: p < 1.2e-05, NES: 1.9) and **antigen processing and presentation of peptide antigen via MHC class I** (FDR p-value: p < 1.2e-05, NES: 1.9). These pathways play key roles in the immune response and neural plasticity, while key genes within the MHC-I protein complex have been implicated in the risk of AD^44,45^. Peptide antigen assembly, processing, and presentation via MHC Class I involves the formation of peptide-MHC complexes, which are crucial for the immune system to recognize and eliminate cells presenting endogenous antigens, including protein aggregates^46^. In AD, these pathways are particularly relevant because dysregulation of immune responses and altered antigen presentation contribute to neuroinflammation and disease progression, especially implicated in microglia^47^. However, given the systemic nature of inflammation and the fundamental role of MHC-I in cellular antigen presentation, these pathways are likely detectable across all cell types, as immune signaling and antigen processing are core processes shared broadly within the cellular environment. Specifically, the aggregation of amyloid-beta impairs antigen presentation by destabilizing the neuronal MHC-I complex^48^. This leads to inadequate immune surveillance and the accumulation of toxic proteins^48^, such as amyloid-beta, as well as dendritic atrophy^49^, which are hallmarks of AD pathology. Consequently, disruptions in these pathways exacerbate neuroinflammation and impair neuronal function, thereby influencing the development and progression of AD.

### Identifying key states enriched for AD heritability enrichment

We next aimed to connect PCA-based states to the genetic etiology of AD through integration with AD GWAS data. We used LD score regression applied to specifically expressed genes (LDSC-SEG)^3^ to detect disease-related states that contained genes enriched for AD heritability (see Methods). Overall, only ATAC-based states showed significant heritability enrichment, while RNA-based states did not (**Fig. 4 A**), consistent with recent work^50^. ResidPCA recovered the most states enriched for AD heritability (ResidPCA: 30 states, Standard PCA: 27 states, Iterative PCA applications: < 9 states) (**Fig. 4 A**), with significant enrichments observed in both early and late PCs (i.e. not just those that explained the most variance) (**Fig. 4 B**). As a benchmark, we compared heritability enrichments to the AD heritability enrichment for genes specific to microglial cells, the established cell type most enriched for AD heritability^51^. Strikingly, ResidPCA captured states with the most significant p-values (minimum p-value < 1.9×10^-8^, FDR corrected) and largest heritability enrichments (maximum enrichment = 4.1), with all PCA methods identifying some states that were more enriched than the microglia annotation (p-value: 0.14) for AD^52^ (**Fig. 4 C**). The enrichment of “later” PCs that explain less variance suggests that subtle single-cell states may still be genetically and biologically meaningful, although often discarded by state based analyses^14,15^. Importantly, these states would have been excluded using a conventional “elbow plot” cutoff rather than the BIC cutoff we employed (**Supplementary Fig. 3**). The heritability analysis supported that using states computed from 20,000 variable genes, as opposed to 3,000 (which is often used in current common practices), did not compromise the robustness of the results; in fact, heritability enrichments were more significant and enriched when using 20,000 variable genes compared to 3,000 (**Fig. 4, Supplementary Fig. 5**).

**Fig 4:**
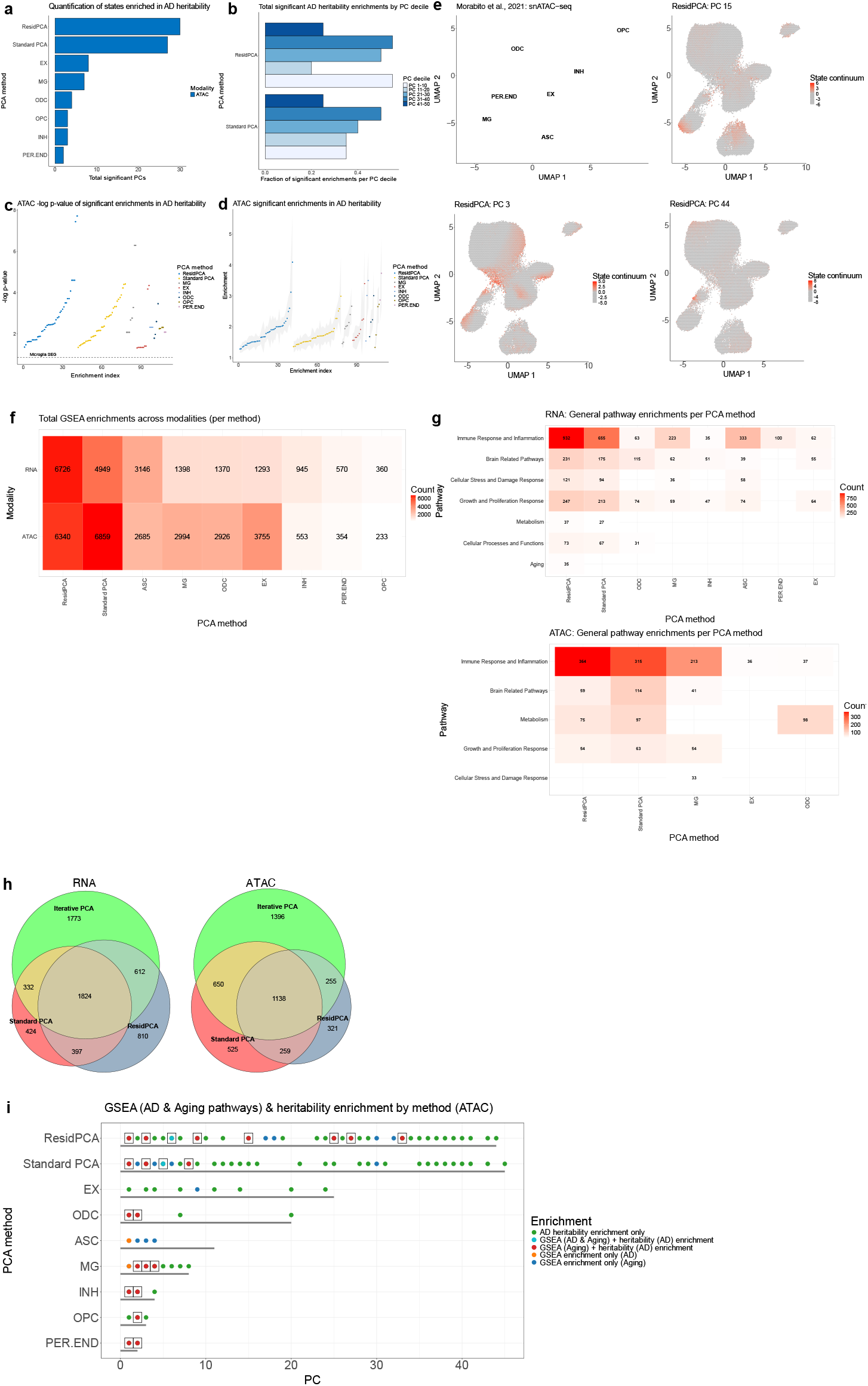
**(a)** Quantification of BIC significant states identified by each PCA-based method (ResidPCA, Standard PCA, and Iterative PCA) enriched for AD heritability in both RNA and ATAC data from Morabito et al. 2021^15^ (FDR p-value < 0.05). **(b)** Fraction of significant ATAC-based states enriched for AD heritability out of the total number of PCs in each decile (10 PCs) in Standard PCA and ResidPCA. **(c)** -log10 p-value or **(d)** heritability enrichment estimate (y-axis) for all significant ATAC-based states enriched for AD heritability across PCA methods, with the respective value for the microglial annotation, the most enriched cell type for AD heritability, also plotted for reference in (**c**). **(e)** Visualization of a subset of ATAC-based states identified by ResidPCA within UMAP space. The top-left plot is color-coded by cell type, while the remaining three plots are colored according to the expression levels of the respective cell embeddings for a given state. **(f)** Heatmap quantifying the number of significant gene set enrichments per method in both RNA and ATAC-based states (FDR p-value < 0.005). **(g)** Heatmaps displaying the number of pathways significantly enriched in gene sets categorized into broader biological pathways per method for both RNA and ATAC-based states. Only broader pathways with more than 25 enrichments in at least one method are included in the plots. **(h)** Euler plots quantifying the number of gene set enrichment across all BIC significant states per PCA method in RNA (left) and ATAC (right) data from Morabito et al. 2021^15^ (FDR p-value < 0.005). Intersections or shared gene set enrichment pathways between Standard PCA, ResidPCA, and/or Iterative PCA are shown. Biological pathways associated with Iterative PCA include the union of gene set enrichments across all seven instances of Iterative PCA applied to each cell type. **(i)** Plot quantifying the colocalization of BIC significant states per method enriched for AD heritability, gene set enrichment for AD, and gene set enrichment for aging in ATAC data. Colocalizations refer to the instances where a state is enriched for AD heritability and the aging gene set and/or the AD gene set. Colocalizations are boxed, and the number of significant states per method is depicted by a gray bar (FDR corrected p < 0.05). Note: (**h**) represents unique pathways, while (**f**) and (**g**) show total pathway counts, including non-unique pathways. Abbreviations: ODC, oligodendrocytes; EX, excitatory neurons; INH, inhibitory neurons; ASC, astrocytes; MG, microglia; PER.END, pericytes and endothelial cells; OPC, oligodendrocyte precursor cells.

Next, we conducted GSEA on ResidPCA-based states most enriched for AD heritability to map disease-based states to their respective biological pathways. The state most significantly enriched for AD heritability was PC15 (AD heritability enrichment: 2.0, FDR p-value: 1.9×10^-8^) (**Fig. 4 C, E**). The enriched pathways in PC15 reveal a novel and recently identified mechanism in AD^16^, spanning the neuron-oligodendrocyte-microglia axis (PC 15 correlation with states identified by Iterative PCA: Excitatory neurons – R^2^: 0.21 , oligodendrocytes – R^2^: 0.14, microglia – R^2^: 0.21, **Fig. 3 D**) . Using GSEA, we found that this state connects early-stage **amyloid production** in neurons and ODCs with later-stage immune response and **microglial activation**, driving AD progression (**Supplementary Table 2**). The state-based annotation, enriched for AD heritability and incorporating the top 200 genes with the most positive loadings, included a genome-wide significant AD locus overlapping **RASGEF1C** (ranked 88/200), suggesting a role for this state in the microglial immune response^53^ (**Supplementary Table 2**). The second most significant ResidPCA-based state for AD heritability, PC27 (AD heritability enrichment: 2.0, FDR p-value: 3.6×10^-8^), relates to AD metabolic dysregulation, enriched in GSEA pathways related to **disruptions in cholesterol** and **steroid hormone metabolism modulated through cytochrome p450**^54,55^ (**Supplementary Table 2**). The most enriched cell types in this state are excitatory neurons and oligodendrocytes (both R^2^: 0.12; though this did not exceed our predefined threshold R^2^ of 0.20 for cell type specificity; **Fig. 3 D**), which are significantly affected by cholesterol and retinol metabolites that influence neuronal plasticity and myelin protection^56–61^. A cell type-specific state identified by ResidPCA is PC44 (AD heritability enrichment: 1.8, FDR p-value: 2.5×10^-5^), which is enriched in microglia (as indicated by GSEA enrichment, though insufficiently powered to reach significant correlation with Iterative PCA microglial states, R2: 0.05) and enriched for AD heritability. This state contrasts with the previously mentioned states that span multiple cell types (**Fig. 4 E, Supplementary Table 2**). It is associated with **microglial activation modulated by CDS+ T cell signaling**, highlighting the interplay between peripheral immune signaling and central nervous system (CNS)-resident microglia in response to AD^62^ (**Supplementary Note 4**).

To assess whether ResidPCA provides additional mechanistic insights compared to Iterative PCA, we aggregated and quantified gene set enrichments across states for each method, rather than interpreting each state individually using GSEA. Overall, ResidPCA identified roughly twice as many significant GSEA enrichments in its RNA (6,726 enrichments) and ATAC (6,340 enrichments) based states compared to Iterative PCA applications to each cell type (RNA: <3,146 enrichments, ATAC: <3,755 enrichments) (**Fig. 4 F**). GSEA enrichments identified by each PCA method were then grouped by pathway into broader categories including inflammatory, brain-related, and metabolic pathways, where the same relationship was recovered (**Fig. 4 G**). In ResidPCA, inflammation-related pathways were the most enriched in RNA-related states (932 enrichments) followed by growth and proliferation response pathways (247 enrichments), and brain-enriched pathways (231 enrichments) (**Fig. 4 G**). Similarly, in ATAC-related states, inflammation was also the most enriched pathway (364 enrichments), followed by metabolism-related pathways (75 enrichments) and brain-related pathways (59 enrichments) (**Fig. 4 G**). To compare the overlap of enrichments between ResidPCA with Iterative PCA applications to each cell type, we analyzed the overlapping and non-overlapping gene set enrichments identified by all Iterative PCA applications versus ResidPCA (**Fig. 4 H**). We found that 56% and 55% of RNA and ATAC enrichments identified by ResidPCA, respectively, did not overlap with Iterative PCA identified enrichments (**Fig. 4 H**). Overall, Iterative PCA, the method typically used to identify states within cell types in sequencing datasets, identified many fewer states and displayed a reduced number of gene set enrichments compared to ResidPCA.

In studies of AD, a key question is whether the pathways driving AD overlap with accelerated age-related processes or represent disease-specific mechanisms independent of general aging phenotypes^63,64^. To address this, we refined our analysis of ResidPCA-based states to distinguish between pathways contributing to AD via aging-related processes and those functioning through disease-specific mechanisms. Using a combination of GSEA and heritability analysis, we evaluated whether the identified states were enriched for AD heritability, gene sets associated with aging, and/or gene sets associated with AD.

Our analysis identified 22 ResidPCA states enriched for AD heritability without significant enrichment for either aging or AD gene sets, suggesting these states are driven by disease-specific pathways independent of aging phenotypes (**Fig. 4 I**). Conversely, PC6 was identified in ResidPCA and showed enrichment for AD heritability (enrichment: 2.1, FDR p-value: 2.5×10^-5^), aging gene sets, and AD gene sets (**Fig. 3D, Fig. 3I and Supplementary Table 2**. Mechanistically, this state reflects pathways linked to both aging and AD-specific processes. This state is predominantly microglial (R2 = 0.23) and encompasses key AD-related pathways, including the **TYROBP microglial signaling network**, which involves the TREM2 receptor that binds amyloid-beta and activates microglial inflammatory responses (**Supplementary Table 2**)^65^. Further analysis of the annotation within this state, enriched for AD heritability and comprising the bottom 200 genes with the most negative loadings in PC6, identified genome-wide significant markers for AD, including **APP** (ranked 195/200), a precursor to amyloid-beta plaques, and **HERC1** (ranked 112/200), associated with protein degradation and inflammation. These findings suggest that PC6 represents a mechanistic intersection of microglial detection of APP and AD-specific inflammatory signaling within the broader age-related pathway of inflammation.

In contrast, we identified seven residPCA states with heritability enrichment and aging gene set enrichment but no AD gene set enrichment (**Fig. 4I**). These states likely contribute to AD through the general acceleration of age-related processes, which are amplified in aging populations but are not exclusive to AD. These pathways represent broad aging mechanisms that predispose individuals to AD as well as other aging-related phenotypes. For example, PC3 is a state spanning three glial cell types—astrocytes (R^2^ = 0.38), microglia (R^2^ = 0.25), and oligodendrocytes (R^2^ = 0.35)—and reflects pathways characteristic of aging rather than AD-specific pathology (**Fig. 3D**). Enriched pathways in PC3 include reduced glial differentiation, impaired axon ensheathment (a key oligodendrocyte function), cholesterol biosynthesis, and protein refolding. These findings suggest that PC3 represents an interactive phenotype where general aging pathways contribute to AD risk without being directly specific to the disease (**Supplementary Note 3**).

Taken together, these findings highlight the distinction between states enriched for AD-specific pathways, such as PC6, which reflect mechanisms directly tied to AD pathology including microglial signaling and amyloid-beta involvement, and those enriched for general aging pathways, such as PC3, which contribute to AD through indirect, age-related mechanisms. By distinguishing between these two enrichment profiles, this analysis advances the understanding of how distinct states drive AD and identifies opportunities for targeted interventions addressing both aging-related vulnerabilities and AD-specific mechanisms.

### Implementation of ResidPCA into efficient computational software package

To facilitate broad adoption of our methodology, we have developed a computational pipeline named the ResidPCA Toolkit. This pipeline efficiently reproduces many of the analyses described in this manuscript, outperforming existing methods in both computational efficiency and resource usage. The ResidPCA Toolkit achieves this by (a) integrating a fast randomized SVD algorithm for principal component calculation and (b) pre-computing a single covariate matrix inversion for expression residuals, avoiding repeated inversions for each target gene.

We evaluated the performance of the ResidPCA Toolkit by comparing it with state-of-the-art methods: Standard PCA implemented with Seurat^66^ and cNMF^12^ implemented with the cNMF package. Our comparison focused on evaluating the maximum memory usage (maximum Resident Set Size [RSS]) and CPU time across a range of simulated count matrix sizes, which varied by both the number of genes and the number of cells in the matrices. Despite handling a more complex task, ResidPCA implemented with the ResidPCA Toolkit required less CPU time than Standard PCA implemented with Seurat and cNMF implemented with the cNMF package (**Fig. 5 A**). Specifically, the average CPU time for ResidPCA using the ResidPCA Toolkit was 0.17 hours (SE: 0.010 In comparison, Standard PCA implemented with Seurat took 0.69 hours (SE: 0.22), and cNMF using the cNMF package required 30.08 hours (SE: 2.70) (**Fig. 5 A**). For a count matrix containing 100,000 cells and 5,000 genes, Standard PCA with Seurat and cNMF with the cNMF package required 9 and 157 times more CPU time, respectively, than ResidPCA using the ResidPCA Toolkit (**Supplementary Fig. 6**). In terms of memory usage, ResidPCA using the ResidPCA Toolkit consumed less memory, with an average maximum RSS of 10.08 GB (SE: 0.029) compared to Standard PCA using Seurat, which used 32.39 GB (SE: 3.22) and ResidPCA using Seurat, which required 47.47 GB (SE: 2.91). cNMF using the cNMF package required slightly less memory, 5.47 GB (SE: 0.087) (**Fig. 5 B**). The key memory difference between cNMF and the ResidPCA Toolkit is that the ResidPCA toolkit saves the outputted embeddings and loadings into memory so that they can be interactively used, however the memory differences are not huge. When comparing the ResidPCA Toolkit to Seurat for performing ResidPCA (an identical task), the ResidPCA Toolkit requires less CPU time and memory (**Fig. 5 A, B**), with its performance scaling linearly with the number of genes (**Supplementary Fig. 6 & Supplementary Note 5**).

**Fig 5:**
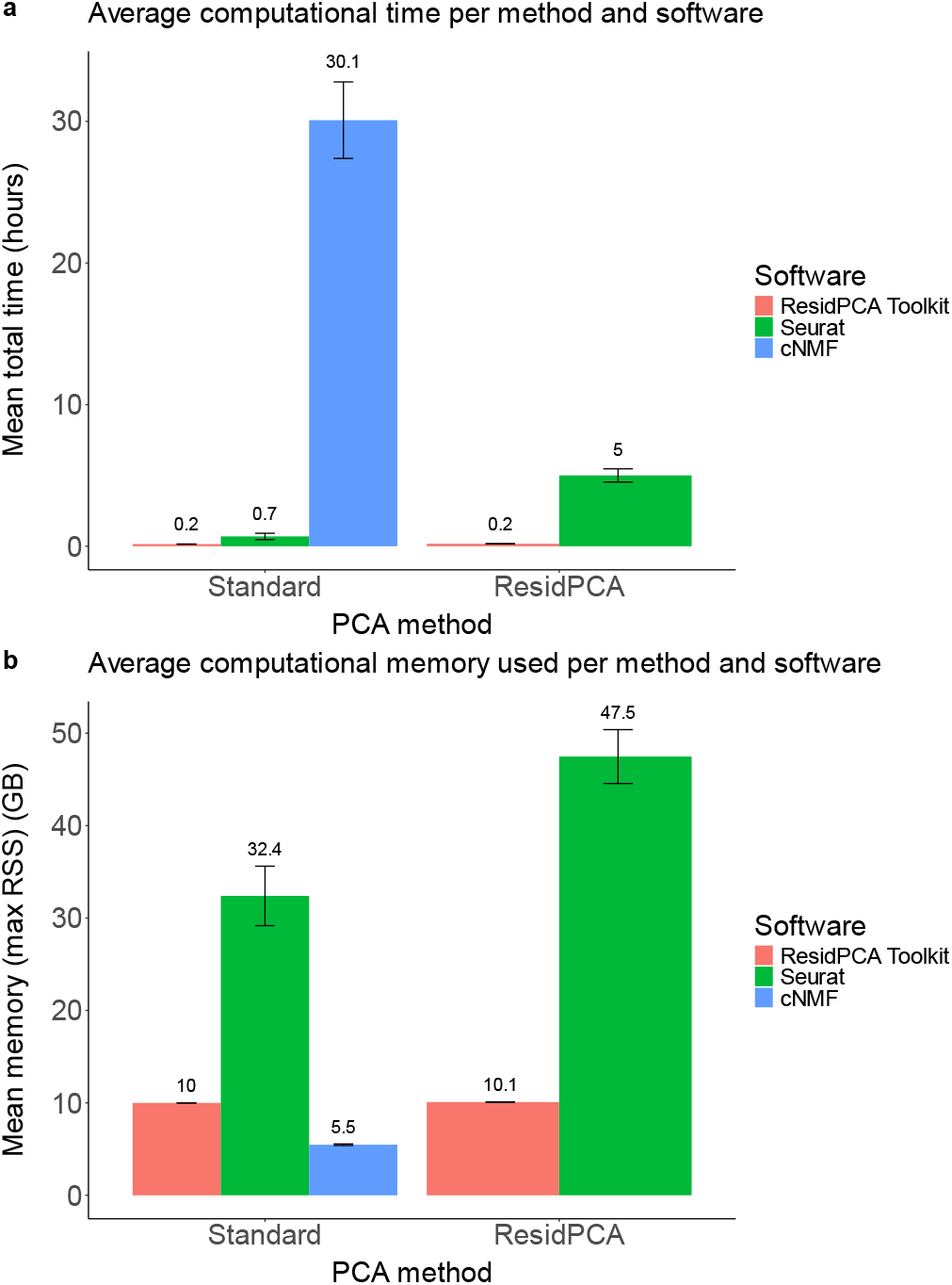
(**a**) Average computational time and (**b**) average computational memory utilized across various input count matrices with differing numbers of cells and genes. The left panels illustrate the average computational time (**a**) and memory (**b**) required by the ResidPCA Toolkit and Seurat to perform Standard PCA, and by the cNMF package to perform cNMF. The right panels depict the average computational time (**a**) and memory (**b**) utilized by the ResidPCA Toolkit and Seurat for executing ResidPCA.

## Discussion

Our study introduces ResidPCA (Residual Principal Component Analysis), a novel method for identifying cellular states in single-cell sequencing data by conditioning out cell type heterogeneity. By analyzing all cells simultaneously, ResidPCA enables the discovery of novel cell states including those that span multiple cell types or are enriched within a single cell type. In contrast, Iterative PCA^14^ and other well-established methods^15,16,19–21^ primarily focus on identifying states within a single predefined cell type. Through extensive simulations and analysis of experimental datasets, ResidPCA excels in identifying a wide spectrum of states ranging from rare to abundant with higher accuracy and sensitivity compared to well-established methods. Notably, ResidPCA was able to identify rare states and those with high cell type heterogeneity in simulation where other methods failed. In real data, ResidPCA greatly outperformed NMF-based methods in detecting light-induced states, particularly in endothelial & smooth muscle and excitatory neurons, the cell types most impacted by light exposure^31^. ATAC-based states produce more enrichment than RNA-based states, consistent with previous analyses^3,50,52^ and ResidPCA consistently revealed more states enriched for heritability than states inferred by other methods or conventional cell type annotations.

The extent to which single cell measurements can provide insights into disease heritability and mechanisms has been an open question^3,5^. In our analyses, state annotations consistently showed higher disease heritability enrichments than cell type annotations, making the identification of cell states beneficial for several reasons^3,5,67^. First, even though disease heritability is typically enriched in genes expressed in relevant cell types or tissues, most of the absolute disease heritability still tends to reside in genes that are broadly expressed across tissues^68^, implicating state-like processes that span many cell types. Second, attributing disease heritability solely to cell types may only tag the disease-related state prevalent in the predominant cell type associated with the disease. However, some cells within this enriched cell type may be “healthy cells” not actively contributing to the disease process. The inclusion of genes from these healthy cells in heritability analyses may weaken the observed heritability enrichment. Finally, cell types can often provide only a high level localization of disease heritability, for example showing that the heritability of neurological conditions is enriched in neurons^3^. By using states as annotations for heritability enrichment, we can pinpoint more specific biological functions driving disease, rather than merely tagging the cell types capable of assuming or not assuming a given disease-related state. This knowledge is crucial for the development of targeted therapeutics that address the specific cellular states driving disease progression, rather than targeting broad cell types.

We make several recommendations for inference of cell states that differ from conventional strategies. First, even after dimensionality reduction methods are applied, inferred states are often visualized using UMAP for face validity and interpretation. However, our simulations showed that two UMAP dimensions often reflect cell state relationships poorly, consistent with prior work^30^, and we caution against interpreting states visually from such a low dimensional representation. Second, while prior methods typically restrict single cell data to a small number of dimensions for downstream clustering (e.g. UMAP for Seurat), we found that single cell datasets contain greater dimensionality in the data than typically considered^10,14,23^. Using the Bayesian Information Criterion (BIC) to determine the number of significant states identified by a given method resulted in many more significant states than using heuristics such as the elbow cutoff. This contrasts with hard clustering, a widely used method^10,14,23^, which assumes that each cell can only occupy one state and that states are exclusively expressed within a single cell type—assumptions that may conflict with established biology^69^. Our observations, showing that many states can be shared across cell types and that individual cells can express multiple states, highlight the limitations of hard clustering and provide insights into how different cell types collaborate to drive disease. Third, single-cell pipelines often restrict to a subset of highly variable genes, with the goal of identifying the most biological variation. However, as cell states are rarer than cell types and capture less overall variation in the data, the ability to detect them can be substantially influenced by the number of genes used in the analysis. Our analyses demonstrate that incorporating additional non-highly variable genes can improve the sensitivity of state detection in some settings but not in others. Defining the precise number or weighting of genes to use for dimensionality reduction in a given experiment thus remains an open question, though we recommend exploratory analyses with a large number of genes.

While ResidPCA is generally more accurate and sensitive to detecting cell states compared to well-established methods, there are some limitations. First, cell state(s) highly correlated with cell type may be poorly detected due to the removal of cell type effects. While we believe the residual cell state component may be particularly informative of biological mechanisms that are independent of cell type, this approach may miss certain cell states or provide a distorted representation of their activity. Second, in modeling cell type expression, discrete cell type labels were used, however continuous or probabilistic cell type assignments can be incorporated which may improve cell type expression estimation, especially for differentiating or difficult to classify cell types. Third, benchmarking methods for cell state inference is hindered by the general lack of well established ground truth cell states. We sought to overcome this limitation by evaluating realistic simulations, experimental data under stimulation, and large-scale data at steady state. However, more refined techniques and standardized datasets of cell states are needed, as have been collected for cell types through single cell atlases^2^. Fourth, ResidPCA does not account for the inherent sparsity in single-cell data, which is driven by technical variations such as stochasticity in cell lysis, reverse transcription efficiency, and molecular sampling during sequencing^70^ as well as biological variation such as transcriptional bursting^71^. While negative binomial or gamma-Poisson models are generally a good fit for this type of data, zero-inflated negative binomial (ZINB) models can sometimes capture sparsity better^72^. This indicates the potential benefit of incorporating a sparsity prior into simulation and state detection.

Alternative approaches for connecting single cell data to heritability include methods like scDRS^73^ and SCAVENGE^74^, which project GWAS data into the single cell space. These methods aim to map genetic associations identified in GWAS to specific cell types and states, providing insights into the cellular context of genetic variants. In contrast, our approach projects single cell annotations onto GWAS heritability, focusing on how single cell data can inform the genetic architecture of diseases. This reverse mapping is effective when there is rich structure in the single-cell data, allowing for precise identification of disease-relevant cellular states. On the other hand, GWAS-based methods may be more advantageous when a large number of causal genes are accurately implicated by GWAS, facilitating the identification of relevant cell types and states. Both approaches offer valuable perspectives, and their effectiveness depends on the complexity and resolution of the underlying data. Additionally, it is worth noting the method described in Lee & Han., 2022^75^, which provides a fast way to perform batch adjusted PCA in single cell data. However, this method does not currently model continuous cell types or covariates, which could be an area for future work.

ResidPCA offers a powerful tool for identifying biologically relevant states and their associated cell types, thereby advancing our understanding of disease mechanisms. Moreover, its ability to leverage all cells enhances statistical power, making it especially valuable in scenarios where the subset of relevant cell types is unknown, rare states are present, or sample size is low. We present a comprehensive computational pipeline and software implementation of ResidPCA, offering researchers a powerful, user-friendly tool to compare PCA methods, uncover novel cell states, and gain new insights into their roles across cell types and complex diseases.

## Methods

### Generative model for cellular states in single-cell data

Let *Z* _*I*×*J*_ represent the log-normalized expression matrix, where *I* denotes the total number of cells and denotes the total number of genes measured in a single-cell experiment. Each entry *z*_*ij*_ in this matrix corresponds to the log-normalized reads per 10,000 (TP10k) for gene *j* within cell *i*. This normalization accounts for differences in sequencing depth across cells, providing a standardized measure of gene expression.

To model *z*_*ij*_, we parameterize it using a Gaussian distribution, which captures the underlying variability and structure in the expression data:

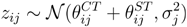

where the mean of *z*_*ij*_ is the sum of two components: the cell type mean expression,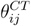 , and the cell state mean expression,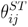. Let 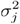 denote the variance for gene *j*.

Next, we define Y_*I*×*J*_ as the observed raw count matrix, which is a transformation of *Z*_*I*×*J*_. In this matrix, each entry *y*_*ij*_ represents the actual number of reads (or counts) for gene *j* in cell *i*. These raw counts are the direct output from sequencing, capturing the number of transcripts in scRNA-seq or the number of peaks in scATAC-seq data.

To model *y*_*ij*_ , we parameterize it using a Gamma-Poisson distribution with respect to the cell type mean expression 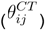 and cell state mean expression 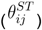:

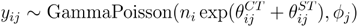

where *n*_*i*_ represents a size factor, such as the total number of unique molecular identifiers per cell and *ϕ*_*j*_ is the overdispersion of gene *j*. The Gamma-Poisson distribution is commonly employed to model raw expression data because it effectively balances simplicity and accuracy, capturing data variability without introducing unnecessary complexity into the statistical model^72^.

### Connecting ResidPCA to single-cell generative model

We now introduce our method Residual Principal Component Analysis (ResidPCA) which leverages log-normalized TP10k data ( *z*_*ij*_) with known cell type labels to estimate the contributions of both cell type and cell state on expression.

The cell type component 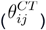 can be expressed as the following inner product:

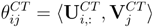

Where 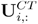 is the cell type indicator vector for observed cell *i* and is the *i* -th row of the cell type indicator matrix 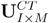 where *I* is the number of cells and *M* is the number of distinct cell types. Additionally,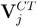 is the normalized mean cell type expression for gene *j* and is the *j* -th column of the cell type mean expression matrix 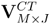 where *j* is the number of genes.

Similarly, the state-driven component 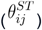 is decomposed into the inner product of the state-driven cell embeddings or principal components (PCs) and the state-driven gene loadings represented below:

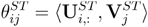

where 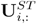, is the *i*-th row of the cell state matrix 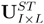, which represent how active each of the *I* cells are in a set of cell states *L*. Separately, 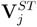 is the *j*-th column of the cell state mean expression matrix 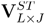, representing how active the set of genes *j* is across a set of *L* states.

Given the above decomposition of 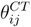 and 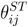, the normalized TP10k expression data (*z*_*ij*_ ) can be redefined as:

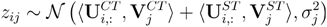

We note that the raw count data *y*_*ij*_ can also be redefined as a function of U_*ST*_, V_*ST*_, U_*CT*_ , and V_*CT*_.

ResidPCA learns the underlying parameters of *Z*_*I*×*J*_ across all genes and cells, namely V^*CT*^ , U^*ST*^ , and V^*ST*^, using a two step procedure. In the first step, V^*CT*^ is learned through linear regression, as U^*CT*^ and Z are known a priori, by minimizing the following negative log-likelihood (i.e. computing the residual):

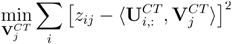

Once V_*CT*_ is learned, maximizing the likelihood of the Gaussian distribution describing *z*_*ij*_ is equivalent to minimizing the following objective function across cells in the second step of our procedure:

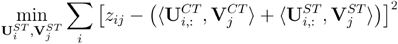

When the above minimization is subject to the orthogonality constraint,

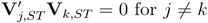

the solution becomes equivalent to using PCA to learn the normalized reconstruction of state-driven expression by minimizing the reconstruction error (or Euclidean distance) between the input data Z and a low-rank, orthogonal representation of the data. The number of states learned, *L*, is a hyperparameter, and its inference is discussed below (see Computing the number of significant components).

### ResidPCA inference model

The goal of ResidPCA is to infer cell states that are independent of cell type. The two state-related parameters inferred by ResidPCA are the normalized cell embeddings (U^*ST*^) and the normalized gene loadings ( V^*ST*^) for a set of *L* states. The state-related cell embeddings quantify the degree to which each cell is involved in a particular state, while the state-related gene loadings measure the contribution of each gene to the set of states. Additionally, ResidPCA infers a cell type-related parameter in an intermediate step: the normalized cell type gene loadings (V^*CT*^) which capture the average expression effect of each gene across the cell types, or the average expression profile of each cell type.

ResidPCA requires two inputs. The first input is the cell type labels, represented by an indicator or probability matrix, U^*CT*^ . We note that U^*CT*^ can also be a probability matrix to account for uncertainty in cell type labels and that other covariates can also be included in the matrix. The second input is the normalized TP10k log-transformed expression matrix, *Z*_*I*×*J*_. This normalization process accounts for sequencing depth, stabilizes variance to reduce skewness, and minimizes the impact of outliers. As a result, the distribution of gene expression data becomes approximately Gaussian with a linear mean-variance relationship. This transformation aligns the data with the assumptions of PCA, including homoscedasticity, linearity, normality, and mean-centering.

ResidPCA is performed in the following steps:

Step 0. <u>Cell Type Label Acquisition:</u> The cell type for each input cell, indexed by *i=*1,….*I*, must be inferred. This can be accomplished using established methodologies such as clustering techniques^76^, marker gene analysis, or integration^77^ with reference datasets. These labels must be obtained prior to performing ResidPCA and are used as input to the analysis.

Step 1. <u>Estimation of Cell Type-Specific Expression Component:</u> For each gene *j*, the cell type-specific expression component, 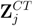, is estimated via linear regression. This regression models the relationship between the cell type indicator (or probability matrix if applicable), U^*CT*^ , and the total normalized expression, Z. The regression coefficients, V^*CT*^, represent the mean expression of gene *j* across different cell types:

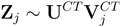

To compute the cell type-specific expression component, 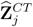, for a given gene *j*, we derive the reconstruction estimates from the regression:

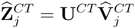

Where 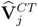 represents the estimated regression coefficients or betas outputted by the regression for a given gene *j*. This step enables precise estimation of gene expression attributable to specific cell types, facilitating further analysis of state-driven expression components.

Note that, instead of computing 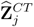 gene by gene, as done in many single-cell analysis tools like Seurat^66^, we compute the matrix 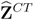 for all genes simultaneously, offering significantly faster performance compared to other implementations.

Step 3. <u>Estimation of State-Driven Expression Component</u>: To estimate the state-driven expression component, 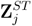, we first subtract the cell type-driven expression component, 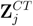 , from the total TP10k normalized expression Z_*j*_. This yields an expression component that is independent of cell type and represents the additive state-driven expression component of gene *j* across all states:

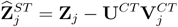

Note that the estimator 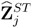 is unbiased with respect to state-driven expression that is independent of cell type. However, it becomes biased if the state-driven expression is dependent or correlated with one or more cell types. This bias arises because any state-driven expression correlated with cell type is regressed out of the total expression matrix.

Step 4. <u>Decomposition of State-Driven Expression Component</u>: To decompose the estimated total state-driven expression component, 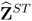 , across all states into distinct states, we apply PCA to the standardized covariance matrix of the estimated state-driven expression component across all genes. Using PCA, which is equivalent to the singular value decomposition (SVD) applied to 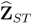 , we identify the state-driven cell embeddings, U^*ST*^, across all cells, the state-related gene loadings, V^*ST*^, across all genes, and the corresponding eigenvalues:

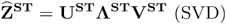

Here, 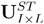 represents the matrix of PCs, illustrating the continuum on which each cell expresses each state. The diagonal matrix 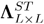 contains singular eigenvalues, ordered by descending variance explained or importance of each PC. The matrix 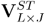 denotes the gene loadings matrix or eigenvectors, indicating the extent to which each gene is expressed within a given state.

Note: The effects of cell type and cell state on 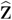 expression on are assumed to be linear.

### Simulation: Synthetic gene expression dataset generation

We generated synthetic gene expression datasets incorporating a set of cell types and a single cell state for simplicity. This was achieved by extending the simulation framework introduced by GlmGamPoi^78^ to simulate concurrent cell state-driven expression alongside cell type-specific expression. Initially, we fit GlmGamPoi to the expression profiles of each cell type separately, using the snRNA-seq data from Morabito et al. (2021)^15^. This allowed us to estimate both the mean cell type expression count vector, 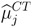, and the overdispersion vector, 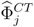, for each gene across the seven cell types in the dataset. Metrics were estimated for 2,527 genes, comprising the union of the top 500 most variable genes in each cell type.

The mean state-driven expression count vector,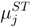 , for all cell types was defined as a function of the mean cell type expression count,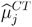 , where:

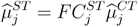

Here, 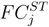 represents the fold change in gene *j* relative to the baseline mean gene expression for each cell type,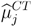 . The fold change, 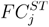, is sampled from:

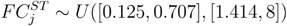

ensuring that:

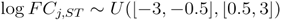

This is consistent with the realistic log *FC* of differentially expressed genes in sequencing data. The transformation reflects that mean cell state-based gene expression is a fold change difference from mean cell type-based expression.

The mean total expression count for a given cell *i* and gene *j* is defined as:

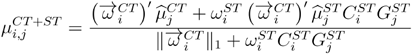

where 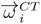 is the indicator vector for a cell’s identity, and 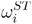 is a scalar representing the continuum on which cell *i* expresses the state, following a standard uniform distribution 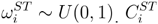 is a binary indicator for state expression in cell *i*, and 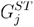 is a binary indicator for state expression in gene *j*.

After generating 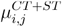 , the mean total expression capturing both cell type and state-driven expression, an expression count for gene *j* in cell *i* is sampled from:

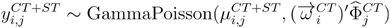

Simulations were conducted to generate count-based gene expression matrices reflecting varying levels of cell type heterogeneity. Datasets included seven cell types to represent low heterogeneity, matching the Morabito et al. dataset^15^, and 100 cell types to represent high heterogeneity. For scenarios with seven cell types, 20,000 cells were simulated, and with 100 cell types, 40,000 cells were simulated. In the latter scenario, the seven fitted cell type parameters from the Morabito dataset^15^ were used alongside 93 *in silica* generated cell type parameters. Mean and overdispersion parameters for these additional cell types were generated by sampling from a uniform distribution bounded by the minimum and maximum parameters of 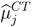 and 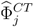 from the real data:

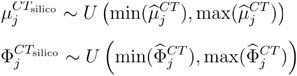

Expression matrices were generated to include states within a single cell type or across all cell types, with states ranging from rare to abundant. This was controlled by varying the number of genes and cells expressing the state, ranging from 0.15 to 0.7 [0.15, 0.3, 0.5, 0.7] of the total genes and cells capable of expressing the state. Cells and genes involved in a given state were randomly sampled, and simulations for each parameter setting were repeated with 10 different seeds.

The simulated datasets were consistent with real snRNA-seq and gene-mapped snATAC-seq datasets in terms of sparsity, mean-variance relationship, and UMAP embedding distribution.

### Computing the number of significant components/latent features

To determine the number of significant components or states recovered by PCA, we used the Bayesian Information Criterion (BIC) method adapted for PCA^79^. The BIC is a statistical criterion used to select the best model by balancing model fit and complexity, penalizing models with more parameters to avoid overfitting. This method is a function of the total number of cells, denoted as *I*, the total number of genes, denoted as *J*, and the sample eigenvalues, λ, obtained from the diagonals of the eigenvalue matrix, Λ. The optimal number of PCs to include in the model is estimated by first computing the BIC value, denoted as BIC_*p*_. Each model *p* includes the first *p* PCs, where *p* ∈ 0,1,…,*J* −1. The dimensionality of the data, or the number of significant principal components (PCs) to retain as significant, is determined by identifying the value of *p* that corresponds to the PC minimizing the BIC value, expressed as 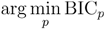. The BIC value is computed as follows:

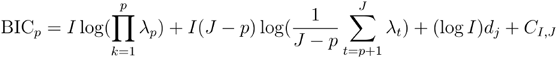

where

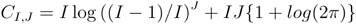

and

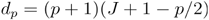

captures the number of independent parameters in the model.

The computation of the sum of eigenvalues greater than *p*, 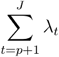, requires evaluating all *J* eigenvalues, where *J* may be as large as 20,000, corresponding to 20,000 total genes. However, to reduce computational and memory demands, only the first 200 eigenvalues or BIC cutoffs were computed, as the significance of PCs was found to be negligible beyond this point. This approach minimizes the computational burden of PCA and the number of BIC cutoff calculations. To avoid computing all eigenvalues, we utilize the fact that the sum of the eigenvalues is equivalent to the trace of the standardized covariance matrix:

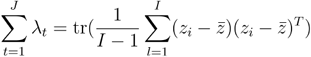

where 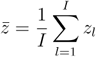 is the average expression across samples. Therefore, the sum of the eigenvalues from *p*+1 to *J* is given by:

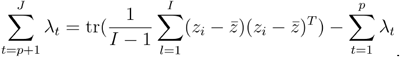

For PCA-based methods, the elbow plot cutoff was also determined by calculating the cumulative variance explained by each successive PC. This process continued until the increase in cumulative variance from adding an additional PC was less than 0.001. The PC at which this criterion was met was selected as the maximum number of PCs to include in the model.

The method for identifying the number of significant components in cNMF was applied to all NMF-based methods^12^. cNMF was executed multiple times, varying the total number of output latent features, denoted as *p*. The optimal number of latent features, *p*, was determined by selecting the model that minimized reconstruction error while maximizing the stability or similarity of results across different runs with the same number of latent dimensions.

### Performance accuracy metrics

We employed two metrics to quantify the accuracy of inferred cell states against a ground truth vector of cell state values.

To assess each method’s ability to infer a single component that maximally captures cell state, we report the maximum *R*^2^ between the vector of true cell state values and the estimated values across all recovered components:

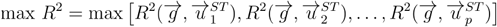

where 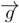 represents the ground truth cell state, and 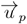 represents a column in the cell embeddings of a given method. The *R*^2^ is computed as follows:

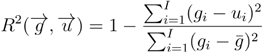

To evaluate each method’s ability to capture overall cell state activity, we report the joint adjusted *R*^2^ from a multivariate regression where the true cell state value vector is the dependent variable and the inferred cell states are the independent variables. The *R*^2^ in this context is computed as:

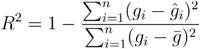

where *ĝ* is the prediction outputted by the multivariate regression. The joint adjusted *R*^2^ is calculated as:

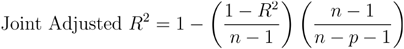

In simulations, *R*^2^ metrics were computed after subsetting to cells putatively expressing the state to distinguish cell state from cell type detection. If the state was expressed in a single cell type, both the embeddings and the ground truth state continuum were subsetted to that specific cell type, and correlations were computed accordingly. Conversely, if the state was expressed across all cell types, the unsubsetted embeddings and state continuum were used.

In the experimental dataset^31^, we computed *R*^2^ after subsetting an embedding or latent feature to a cell type of interest and calculated the corresponding *R*^2^ for each cell type. Additionally, we used the duration of light exposure as the ground truth cell state vector for each state. Since the ground truth continuum of the cell state is unknown in the experimental dataset, we assumed a linear relationship between light exposure and latent cell state.

In simulation, metrics for determining the number of components for dimensionality reduction methods were not used, as the optimal number of PCs or factors was known. In simulations with seven cell types plus one state, 10 PCs were inferred; in simulations with 100 cell types plus one state, 105 PCs were inferred. In the experimental dataset, metrics for determining the number of components for dimensionality reduction methods was used (as described in the above section).

### Cell type annotation and data preprocessing

The single-cell datasets from Hrvatin et al.^31^ and Morabito et al.^15^ were re-processed using Scanpy^80^. This involved filtering out cells with high mitochondrial read counts, removing gene outliers to maintain a linear mean-variance relationship, and ensuring that each gene and cell contained a substantial number of reads. Both datasets were originally annotated with cell type labels as part of their respective studies.

For snATAC-seq datasets, peak data were condensed into a gene activity matrix using the “CreateGeneActivityMatrix” function in Signac within Seurat^66^. This function associates peaks with their nearest gene if the peak is located within the gene body or up to 2kb upstream. The output is a gene-by-cell matrix that can undergo quality control as previously described, owing to its transformed dimensionality from cells by peaks to cells by genes.

The count-based gene-by-cell matrices were then log-normalized and scaled to ensure a total expression of 10,000 reads per cell (TP10k normalization). Subsequently, the data were standardized to a mean of 0 and a variance of 1 for each gene. Variable genes were selected using the ‘highly_variable_genes’ function in Scanpy^80^.

In the Hrvatin et al., 2018^31^ dataset, we tested both 3,000 and 20,000 variable genes as inputs for each method to assess whether a higher or lower gene threshold resulted in better performance. In the Morabito et al., 2021^15^ analysis, we used the top 20,000 most variable genes to minimize ad hoc thresholding and capture all genes in the genome. To determine the number of significant cellular states, we employed the BIC adapted for PCA^79^.

A downsampling analysis of the scRNA-seq expression matrix was conducted using data from Hrvatin et al., 2018^31^ to evaluate the sensitivity of each dimensionality reduction method in detecting light-induced states with fewer samples. The entire quality-controlled dataset was downsampled to 35,000, 25,000, 15,000, 10,000, and 5,000 cells, respectively, and subsequently used as input for each method. Cells were randomly selected for downsampling, and this process was repeated with five different random seed iterations. The BIC method was employed to determine the number of significant states for PCA-based methods, while Error-Stability Optimization was used for NMF-based methods.

In the Morabito et al., 2021^15^ dataset, we quantified the fraction of variation explained by cell type using the following approach. First, we regressed out all covariates except for cell type from the snATAC or snRNA-seq matrix. The resulting cells-by-genes matrix was then standardized. Next, each column of this standardized matrix, representing gene-wise data, was regressed on cell type. We calculated the predicted cell type gene expression matrix by multiplying the cell type by the recovered beta coefficients. Finally, the average variance of this predicted matrix was computed and reported as the average variance explained by cell type.

### Heritability enrichment analysis with LDSC-SEG

We investigated the AD heritability using established GWAS associations from the latest AD GWAS by Bellenguez et al. (2022)^34^. We employed LDSC-SEG^3^, a methodology designed to partition polygenic trait heritability based on functional annotations, to quantify the contribution of our identified states to AD heritability. To define a state annotation, we selected the top and bottom 200 genes within each PC-derived state as separate annotations for input into LDSC-SEG. The top and bottom 200 genes were selected because PCA components are mean-centered, and directionality is not known. To identify differentially expressed genes associated with specific cell types, we used the Scanpy^80^ ‘rank_genes_groups_df’ function. The top 200 ranked differentially expressed genes from each cell type were also used as annotations for input into LDSC-SEG. We retained annotations with significant enrichment scores at an FDR-corrected p-value threshold, categorizing states as significantly enriched for AD heritability if they met this criterion.

### Characterizing cell states

To determine whether a given cell type was involved in a cell state, we calculated the maximum squared correlation between each state identified by ResidPCA or Standard PCA and the respective cell type-specific states identified by Iterative PCA methods. For example, to assess whether a state derived from ResidPCA was expressed in excitatory neurons, the maximum correlation was calculated between the ResidPCA state and all significant Iterative PCA-derived states identified in neurons. The resulting distribution of squared correlations was graphed per cell type to determine a correlation cutoff at the saddle point of the bimodal distribution.

States identified by ResidPCA exceeding this cutoff were considered to be expressed in the corresponding cell type. States were labeled as cell-type specific or agnostic based on the squared correlation between each state and cell-type specific states identified by Iterative PCA methods. Cell type agnostic states are those that ResidPCA can detect, whereas state-of-the-art methods like Iterative PCA lack this capability, underscoring ResidPCA’s superior ability to identify a variety of states.

The GSEApy package^81^ in Python was used to compute gene set enrichments. Five collections of gene sets from the Human MSigDB were used: the human hallmark gene sets, human positional gene sets, human curated gene sets, human regulatory target gene sets, and human ontology gene sets. For mouse analyses on the Hrvatin et al. (2018)^31^ study, gene ontology enrichments from the study were used, along with two gene sets from Mouse MSigDB^82^ (mouse-ortholog hallmark gene sets and mouse curated gene sets), and a recently curated dataset containing gene sets related to the mouse nervous system. Additionally, we used a recently published gene set specific to beta amyloid production^16^.

## Acknowledgements

We extend our sincere gratitude to B. Tolooshams, A. Lin, D. Ba, M. Theodosis, and D. Yao for their valuable insights during discussions on computational methods and single-cell data. We also thank Y. Ding for their creative input in naming the method. S.C. acknowledges support from the National Science Foundation Graduate Research Fellowship Program.

## Author contributions

A.G. and S.C. conceptualized the study, designed the methodology, and led the statistical analysis. All authors assisted in computational analyses, provided revisions to the manuscript and contributed to the final draft. All authors read and approved the final manuscript.

