## Supplementary Figures and Notes for "Discovery of disease-associated cellular states using ResidPCA in single-cell RNA and ATAC sequencing data"

### Supplementary Figure 1: Simulation results across parameter space in 7 cell types.

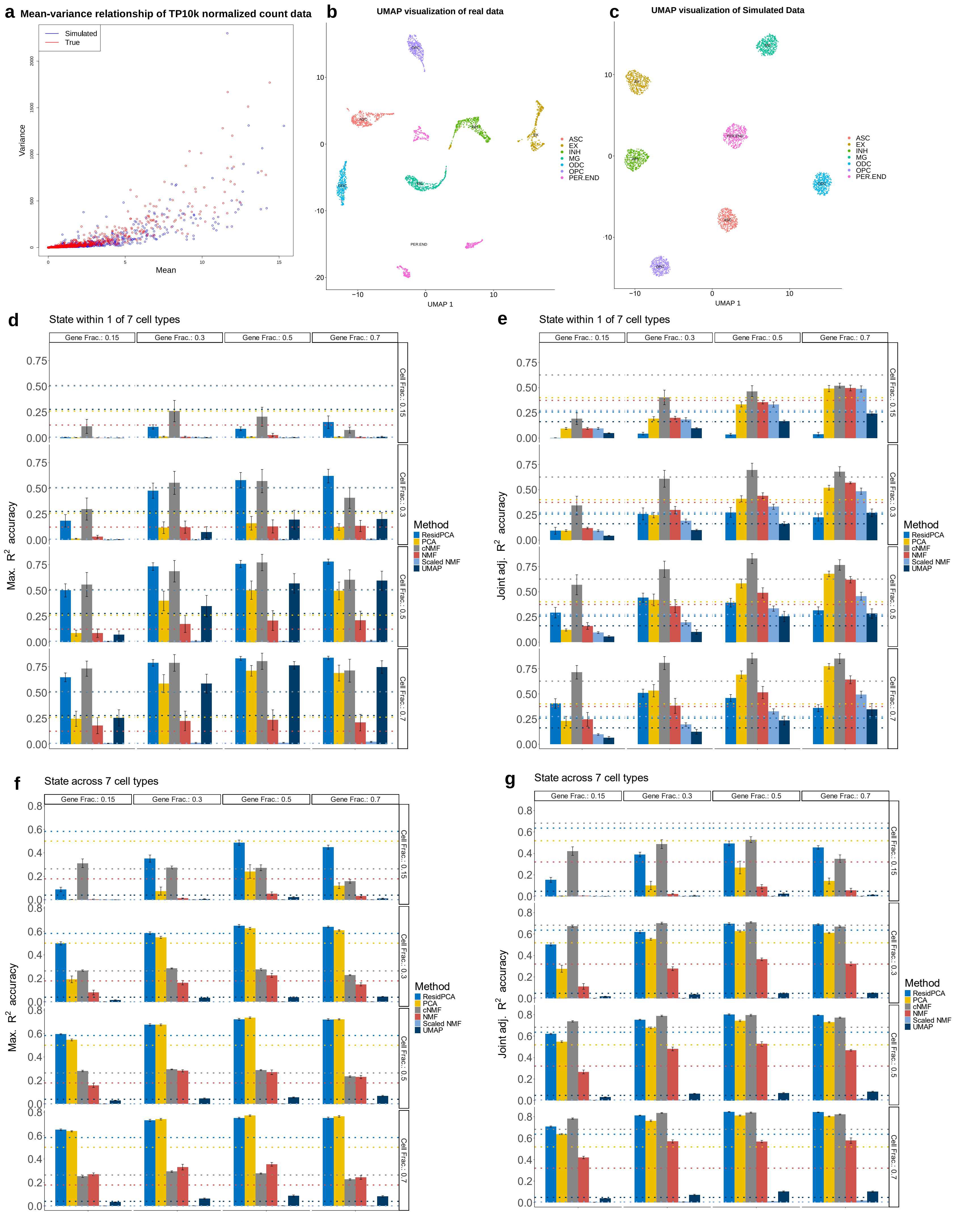

**Supplementary Figure 2: Classification of light derived states and comparative analysis using 20,000 variable genes vs. 3,000 variable genes.**

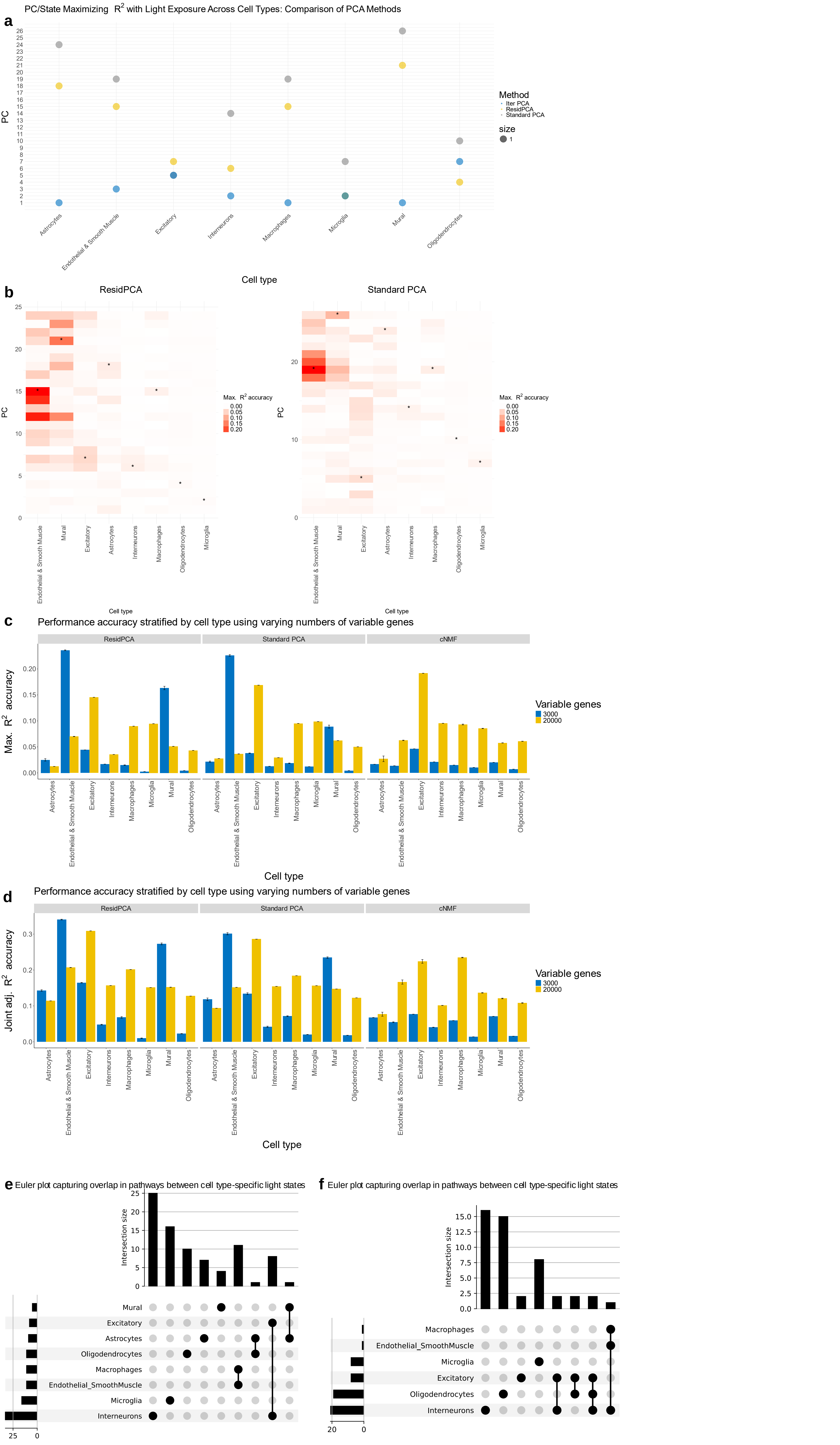

**Supplementary Table 1: GSEA enriched pathways relating to light exposure.**

| GSEA Term | Cell Type | NES | FDR q-val | Tag % | Gene % |
|---|---|---|---|---|---|
| Gray_2014_AllStressConditions_HC | Interneurons | 1.767 | 0.004 | 23/93 | 6.33% |
| BUYTAERT_PHOTODYNAMIC_THERAPY_STRESS_UP | Interneurons | 1.836 | 0.002 | 27/132 | 5.40% |
| Gray_2014_ChronicRestraintAndForcedSwimStress_Upregulated_HC | Excitatory | 1.910 | 0.0 | 12/31 | 4.27% |
| FAN_EMBRYONIC_CTX_MICROGLIA_1 | Microglia | 1.746 | 0.0 | 39/52 | 8.57% |
| Gray_2014_ChronicRestraintAndForcedSwimStress_Upregulated_HC | Endothelial_Smooth Muscle/Macrophages | -1.853 | 0.006 | 11/31 | 3.23% |
| Park_2011_Coexpression_Hippocampus_Mouse_greenyellow | Mural | 2.016 | 0.0 | 15/32 | 8.40% |

**Supplementary Figure 3: Elbow plot vs. BIC & State recovery using 3,000 variable genes.**

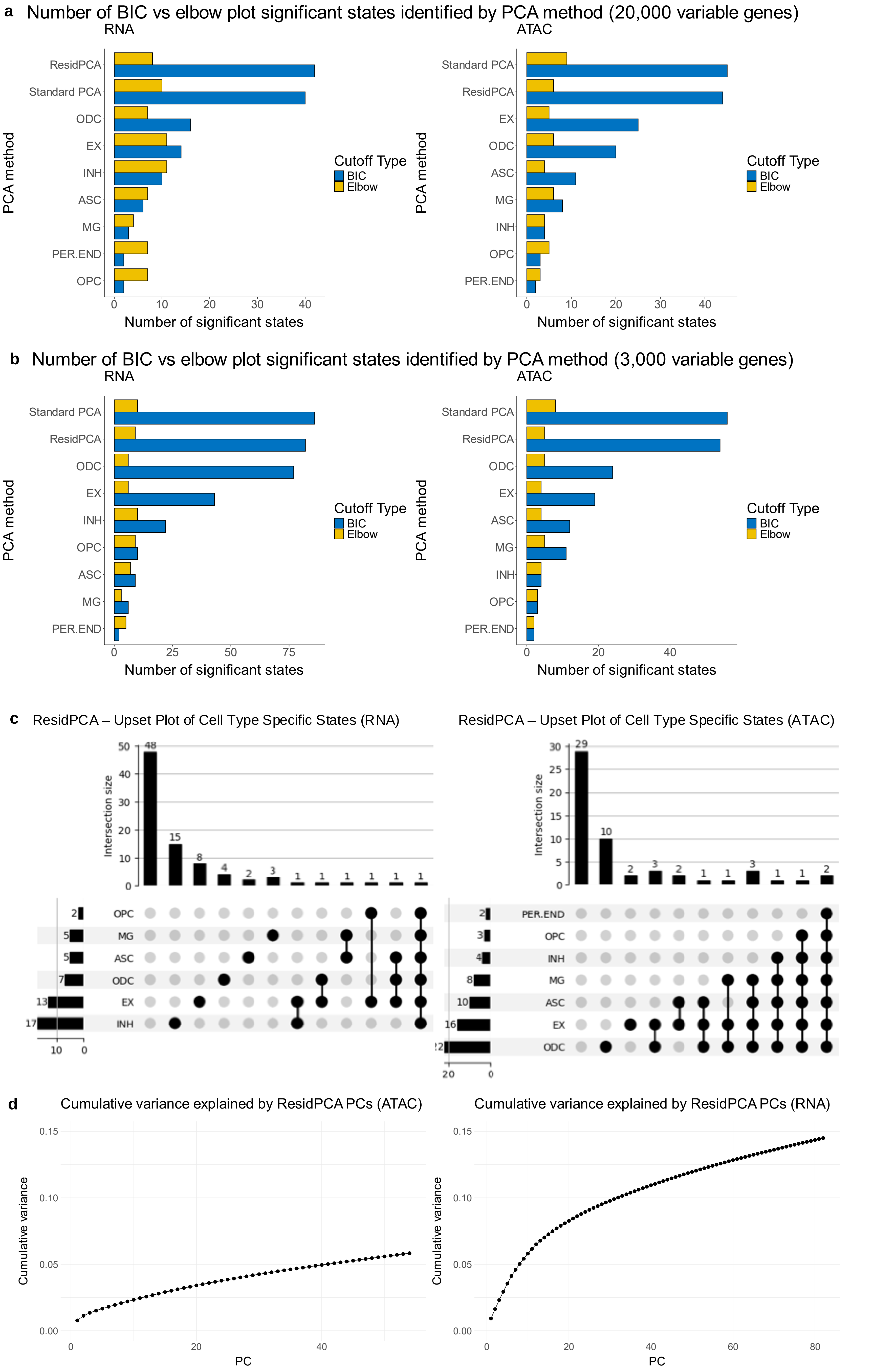

**Supplementary Figure 4: Selecting cutoff for cNMF according to Kotliar et al. recommendation.**

a

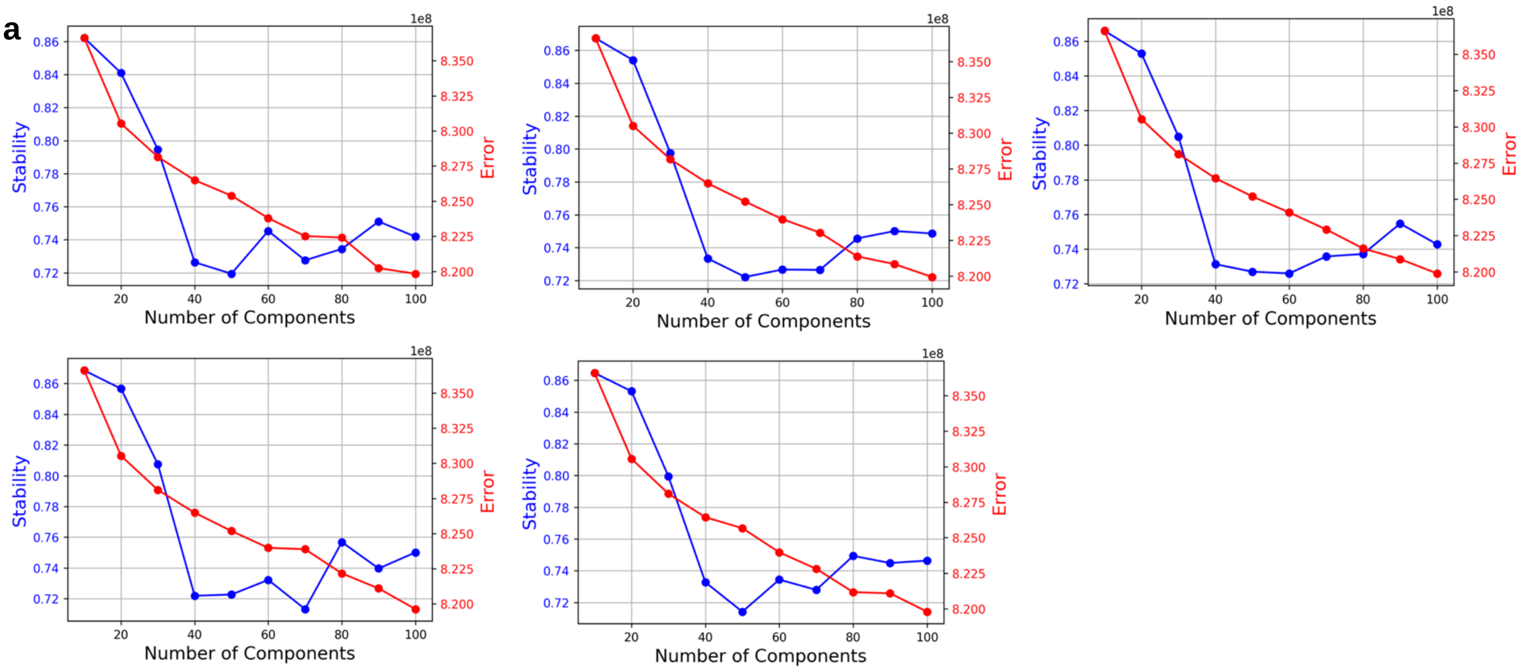


b

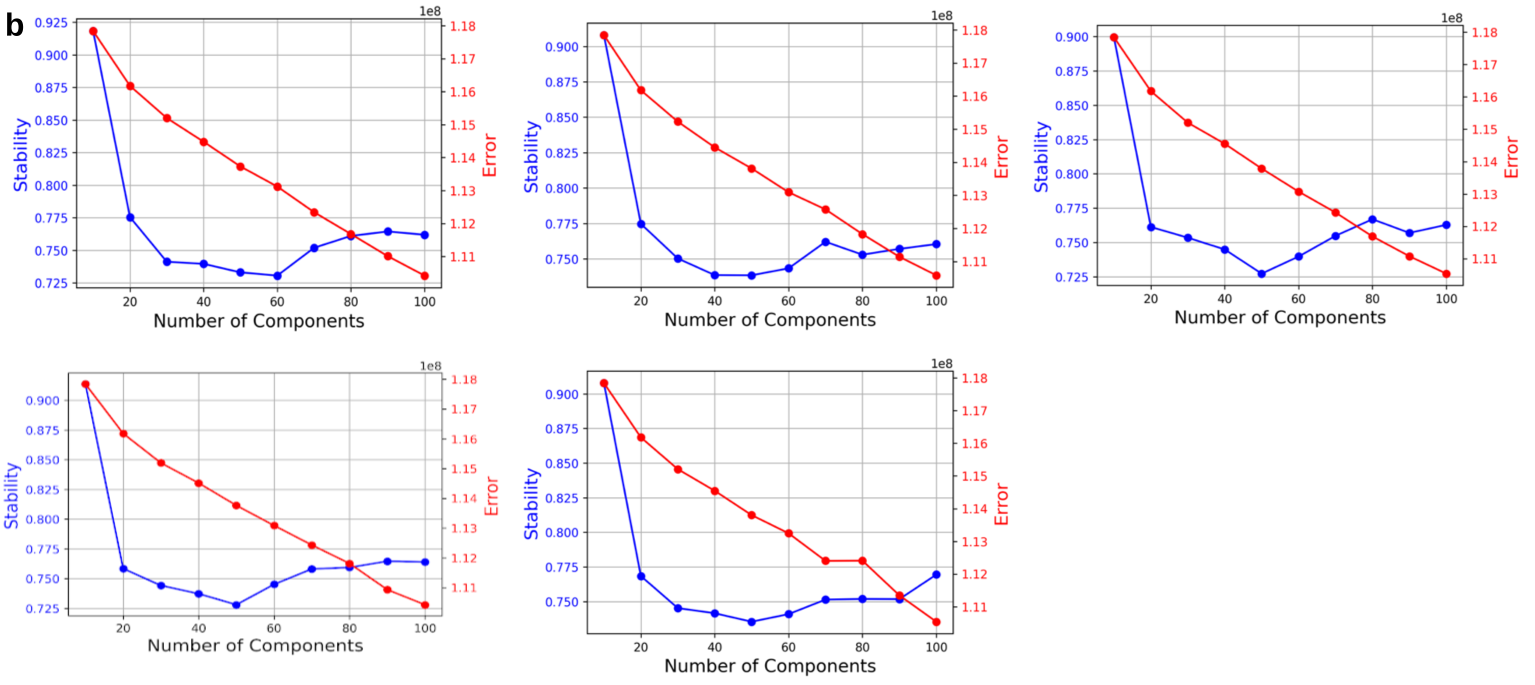


c

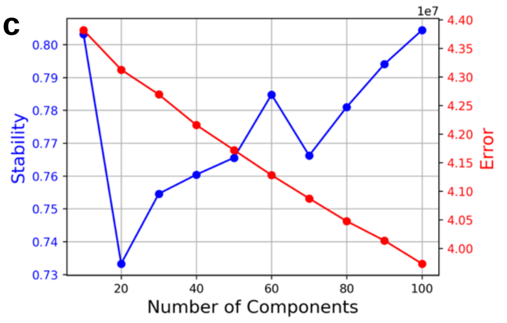


d

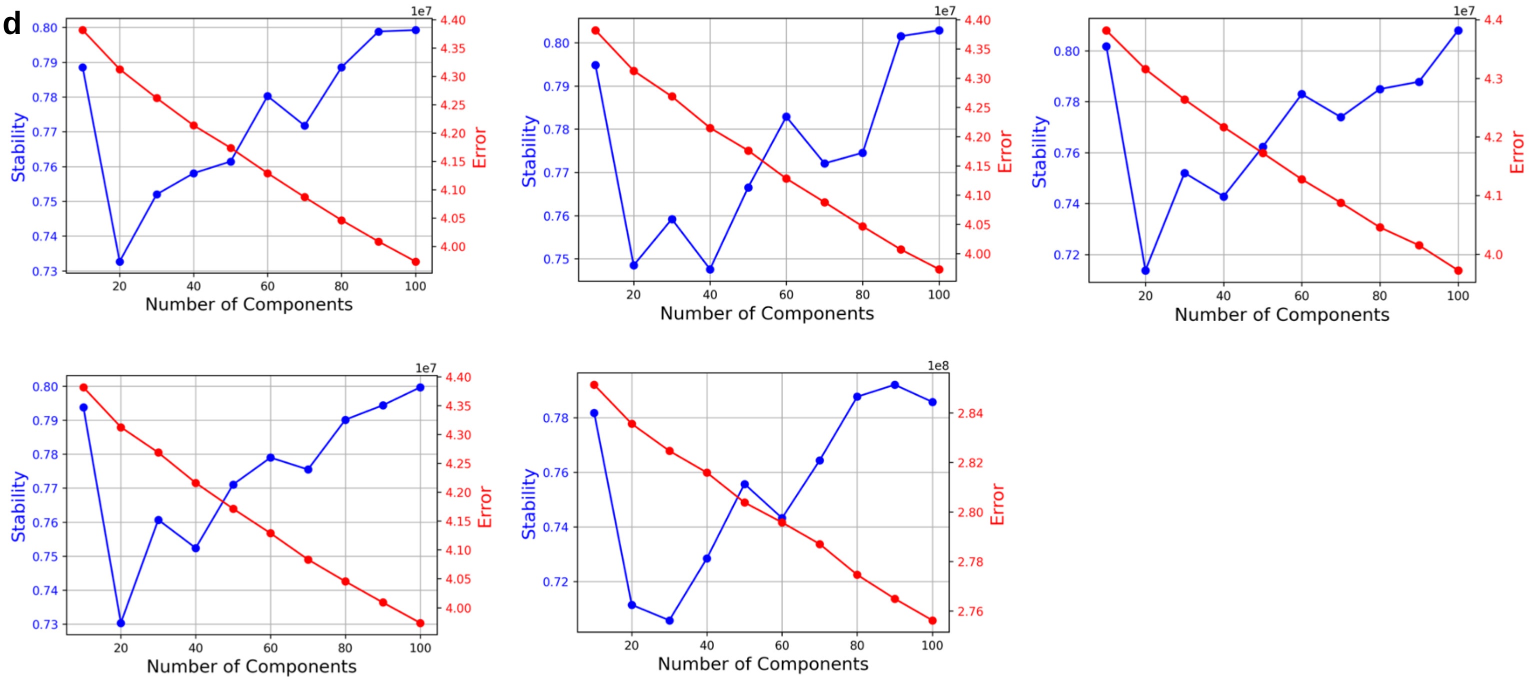

### Supplementary Figure 5: States classified by AD heritability and gene set enrichment using 3,000 variable genes.

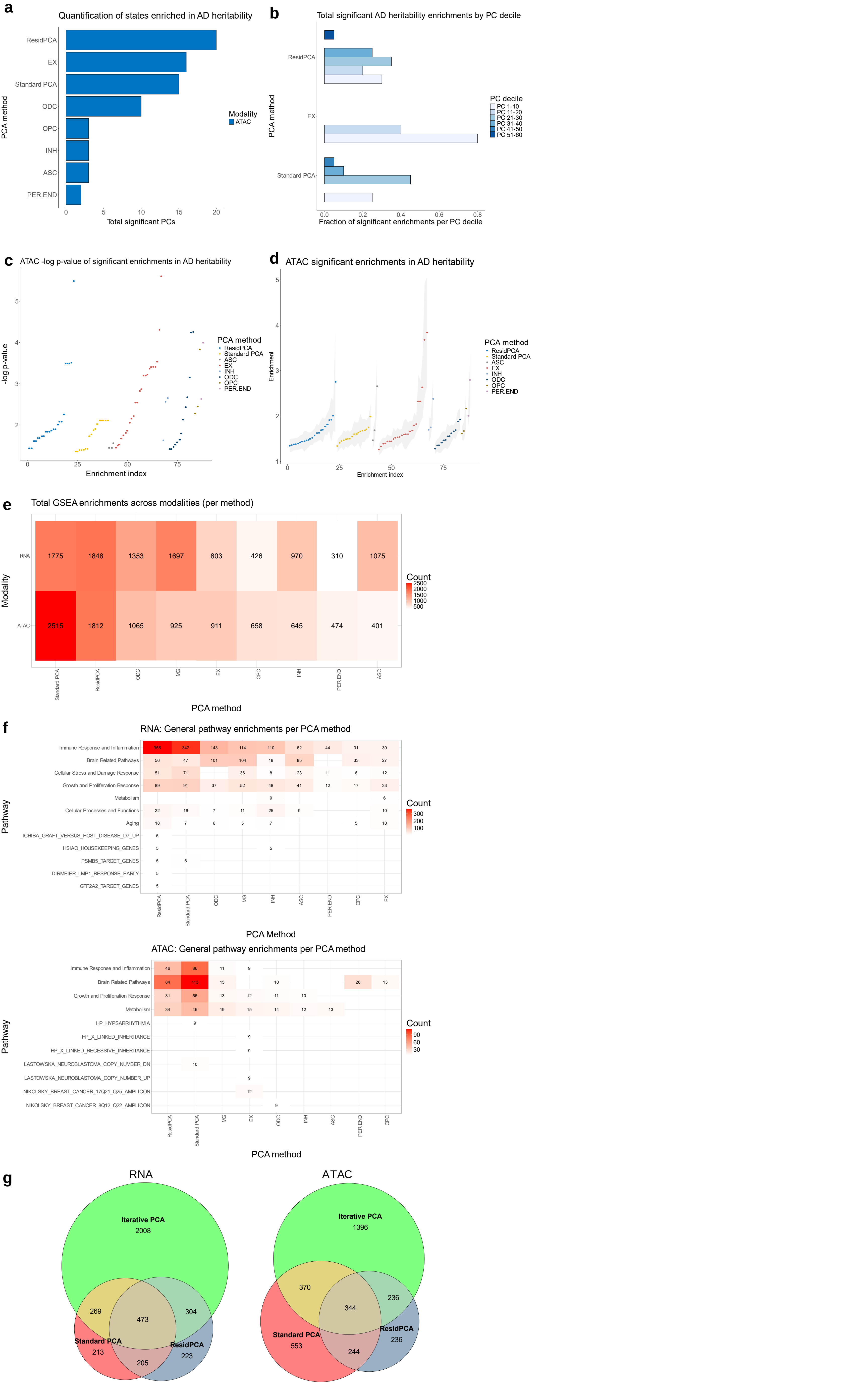

#### Supplementary Figure 6: Comparative Analysis of CPU Time and Memory Usage Across Methods and Packages Under Varying Input Count Matrix Sizes.

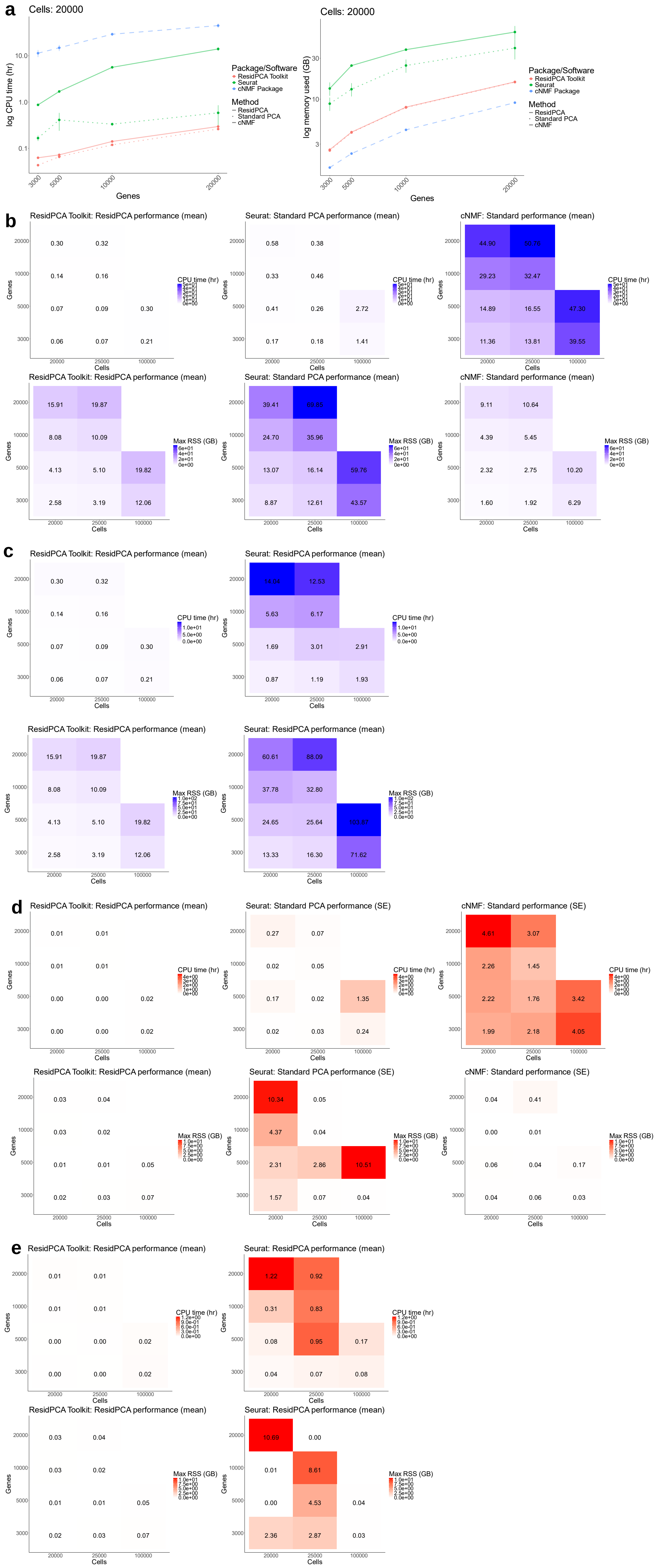

**Supplementary Table 2: Key GSEA enrichments of ResidPCA-identified states.**

| GSEA Term | ResidPCA State | NES | FDR q-val |
|---|---|---|---|
| Antigen processing and presentation of exogenous peptide antigen | PC15 | 2.509 | p < 1.2e-05 |
| Microglia pathogen phagocytosis pathway | PC15 | 2.507 | p < 1.2e-05 |
| Hallmark interferon alpha response | PC15 | 1.934 | 0.004 |
| Regulation of inflammatory response to antigenic stimulus | PC15, | 1.933 | 0.004 |
| Regulation of CD4 positive alpha beta T cell differentiation | PC15 | 1.932 | 0.004 |
| Beta amyloid production | PC15 | 1.550 | 0.009 |
| Drug metabolism cytochrome P450 | PC27 | -3.414 | 1.2e-05 |
| Steroid hormone biosynthesis | PC27 | -3.207 | p < 1.2e-05 |
| Retinol metabolism | PC27 | -3.134 | p < 1.2e-05 |
| Pentose and glucuronate interconversions | PC27 | -2.929 | p < 1.2e-0 |
| Modulators of TCR signaling and T cell activation | PC44 | 2.159 | 0.001 |
| CD8 TCR downstream pathway | PC44 | 2.145 | 0.002 |
| Differentiating T lymphocyte | PC44 | 2.162 | 0.002 |
| AD | PC6 | 1.771 | 0.013 |
| Aging | PC6 | 2.912 | p < 1.2e-05 |
| MIR221* | PC6 | -3.062 | p < 1.2e-05 |
| MIR4776_3P* | PC6 | -3.048 | p < 1.2e-05 |
| MIR362_3P* | PC6 | -3.003 | p < 1.2e-05 |
| MIR1322 | PC6 | -3.301 | p < 1.2e-05 |
| MIR10393_3P | PC6 | -3.287 | p < 1.2e-05 |
| MIR767_3P | PC6 | -3.280 | p < 1.2e-05 |
| MIR6824_5P | PC6 | -3.269 | p < 1.2e-05 |
| MIR4718 | PC6 | -3.267 | p < 1.2e-05 |
| MIR759 | PC6 | -3.268 | p < 1.2e-05 |
| MIR6801_5P | PC6 | -3.266 | p < 1.2e-05 |
| MIR6793_3P | PC6 | -3.233 | p < 1.2e-05 |
| TYROBP causal network in microglia | PC6 | 2.896 | p < 1.2e-05 |
| TNFA signaling via NFKB | PC6 | 2.944 | p < 1.2e-05 |
| Aging | PC3 | -3.365 | p < 1.2e-05 |
| Oligodendrocyte markers | PC3 | -4.151 | p < 1.2e-05 |
| Cholesterol biosynthesis | PC3 | -2.928 | p < 1.2e-05 |
| Axon ensheathment in CNS | PC3 | -2.811 | p < 1.2e-05 |
| Protein refolding | PC3 | -2.589 | p < 1.2e-05 |
| Regulation of glial cell differentiation | PC3 | -2.272 | 0.004 |

***** indicates microRNAs previously established to be associated with AD.

**Supp Fig. 1 description:** (**a**) Mean-variance plot illustrating the relationship between gene expression in the simulated dataset (Simulated) and the true data (True) obtained from the count matrix in Hrvatin et al., 2018[31]. The true data was used to fit the underlying Gamma-Poisson parameters from which the simulated data was generated. (**b**) UMAP visualization of the real data derived from the Hrvatin et al. 2018 scRNA-seq count matrix (True), compared with (**c**) the UMAP plot of the simulated count matrix generated by fitting the GlmGamPoi model to the true data. Note: No cell states were simulated in these plots. (**d, e**) Performance of methods in simulation scenarios with a single state present within one of seven cell types, and (**f, g**) scenarios where a single state spans all seven cell types. In these scenarios, both the fraction of cells expressing the state (y-axis) and the fraction of genes expressing the state (x-axis) were varied. Ten simulation replicates were averaged for each scenario. The maximum $R^2$ accuracy for a single state was calculated in (**d**) and (**f**), while the joint adjusted $R^2$ accuracy across all significant states was computed in (**e**) and (**g**).

**Supplementary Fig. 2 description:** (**a**) Dot plot illustrating the BIC-significant PC or state (y-axis) that maximized the $R^2$ between the recovered PC embedding and light exposure duration for each cell type (x-axis). The results compare Standard PCA, ResidPCA, and separate Iterative PCA runs per cell type. (**b**) Heatmaps depicting the $R^2$ correlation between each PC/significant state embedding (y-axis) and light exposure duration, stratified by cell type (x-axis). Asterisks (*) denote the significant state/PC with the highest $R^2$ correlation per cell type for ResidPCA (left) and Standard PCA (right). (**c, d**) Performance comparison of using 3,000 versus 20,000 variable genes as input for each method, shown by the maximum $R^2$ (**c**) and joint adjusted $R^2$ (**d**) between light exposure duration and the significant states identified by each method, stratified by cell type on the x-axis. Top methods are compared including ResidPCA, Standard PCA, and cNMF. (**e**) Upset plot demonstrating the overlap and uniqueness of GSEA pathways within each cell type-specific ResidPCA state that maximized the correlation between light exposure duration and state embedding for each cell type (FDR p-value < 0.01). (**f**) Similar Upset plot focusing on a curated set of mouse brain-related pathways (FDR p-value < 0.01). Except for panels (**c**) and (**d**), analyses are conducted using 3,000 variable genes.

**Supplementary Table 1 description:** Table of relevant GSEA terms and corresponding statistics for pathways associated with light exposure or conditional stress. The GSEA pathways are limited to terms enriched in each ResidPCA state that maximized the correlation between light exposure duration and the state-driven cell embedding, specific to the cell type of interest (FDR p-value < 0.01). Gene set enrichments are performed on states identified using 3,000 variable genes.

**Supplementary Fig. 3 description:** The comparison of the number of significant states identified by each PCA-based method is shown for single-cell matrices with either 20,000 variable genes (**a**) or 3,000 variable genes (**b**), highlighting the results obtained using the BIC method versus the Elbow plot method. The analysis was conducted on snRNA-seq (left) and snATAC-seq (right) data from Morabito et al., 2021[15]. Methods named after specific cell types correspond to Iterative PCA applied to that cell type. (**c**) Frequency of ResidPCA recovered states spanning one, multiple, all, or no cell types in RNA (left) and ATAC (right) data when 3,000 variable genes were used as input rather than 20,000 variable genes (in **Fig. 3**). (**d**) Cumulative variance plot illustrating the proportion of total variance captured by PCs as more PCs are included, up to the BIC cutoff. The plots show the cumulative variance explained for ResidPCA applied to RNA (left) and ATAC (right) datasets, respectively. Both plots are based on 3,000 variable genes. Abbreviations: ODC, oligodendrocytes; EX, excitatory neurons; INH, inhibitory neurons; ASC, astrocytes; MG, microglia; PER.END, pericytes and endothelial cells; OPC, oligodendrocyte precursor cells.

**Supplementary Fig. 4 description:** Stability vs. error plots generated by the cNMF package from Kotliar et al., 2019[12] using different seeds on the Hrvatin et al. 2021[31] dataset. The analyses were conducted with (**a**) 20,000 variable genes across all QC'd cells, (**b**) 3,000 variable genes across all QC'd cells, (**c**) 20,000 variable genes limited to cells classified as interneurons and excitatory neurons, and (**d**) 3,000 variable genes limited to cells classified as interneurons and excitatory neurons.

**Supplementary Fig. 5 description:** **(a)** Quantification of BIC significant states identified by each PCA-based method (ResidPCA, Standard PCA, and Iterative PCA) enriched for AD heritability in both RNA and ATAC data from Morabito et al., 2021[15]. **(b)** Fraction of significant states enriched for AD heritability out of the total number of PCs in each decile (10 PCs) in Standard and ResidPCA. -log10 p-value **(c)** or heritability enrichment estimate **(d)** for all significant states enriched for AD heritability across PCA methods. **(e)** Heatmap quantifying the number of significant gene set enrichments per method in both RNA and ATAC-based states (FDR p-value < 0.005). **(f)** Heatmaps displaying the number of pathways significantly enriched in gene sets categorized into broader biological pathways per method for both BIC significant RNA and ATAC-based states. Only broader pathways with more than 25 enrichments in at least one method are included in the plots. **(g)** Euler plots quantifying the number of gene set enrichment across all BIC significant states per PCA method in RNA (left) and ATAC (right) data from Morabito et al. 2021[15] (FDR p-value < 0.005). Intersections or shared gene set enrichment pathways between Standard PCA, ResidPCA, and/or Iterative PCA are shown. Biological pathways associated with Iterative PCA include the union of gene set enrichments across all seven instances of Iterative PCA applied to each cell type. Note: This figure was generated using 3,000 variable genes as input for each PCA-based method, in contrast to the 20,000 genes used in Figure 4. Abbreviations: ODC, oligodendrocytes; EX, excitatory neurons; INH, inhibitory neurons; ASC, astrocytes; MG, microglia; PER.END, pericytes and endothelial cells; OPC, oligodendrocyte precursor cells.

**Supplementary Fig. 6 description:** (**a**) A plot illustrating the average performance of each method and package as a function of matrix input size, achieved by varying the number of genes in the input count matrix while maintaining a constant number of cells. The resulting CPU time (left) and memory usage (maximum resident set size [RSS], in GB) (right) for each method. Each point is the average across five replicates. (**b,c**) Plot of average CPU time (left) and memory usage (right) across all replicates and runs, as a function of input count matrix size. The input count matrix size was varied using a grid of parameters, adjusting both the number of cells and the number of genes. (**b**) CPU time (top) and memory usage (bottom) are compared for ResidPCA implemented with the ResidPCA Toolkit, Standard PCA implemented the Seurat package, and cNMF implemented with the cNMF package. The metrics are shown for count matrices with varying numbers of cells and genes. (**c**) The comparison of CPU time (top) and memory usage (bottom) between ResidPCA implemented with the ResidPCA Toolkit and implemented with Seurat. (**d**) Similar to plot (**b**), but instead of showing the mean, the standard error is plotted across replicates. (**e**) Similar to plot (**c**), but instead of showing the mean, the standard error is plotted across replicates.

**Supplementary Table 2:** Key GSEA enrichments of ResidPCA-identified states.

#### Supplementary notes

##### Supplementary Note 1, Fig. 1: Distinct Light-Induced States Across Cell Types Revealed by ResidPCA

We investigated whether ResidPCA identified a single light-induced state universally expressed across all cell types or distinct light-induced states specific to each cell type. To assess the similarity of light-induced states across each cell type within a specific method, we calculated the pairwise cross-correlation between the states that showed the highest correlation with the duration of light exposure for each cell type. Our analysis revealed that each cell type's light exposure duration correlated maximally with a distinct PC, except for endothelial & smooth muscle cells and macrophages.(**Supplementary Fig. 2**). This suggests that while most cell types exhibit unique light-induced states reflected in distinct states/principal components (PCs), endothelial & smooth muscle cells and macrophages may respond to light exposure in a similar manner, indicating potential shared mechanisms or pathways in their light response. Previous research has shown that macrophages residing in the vascular wall closely interact with endothelial cells, facilitating the process of angiogenesis[83]. Additionally, ResidPCA distinguishes light-induced states more clearly between mural cells and endothelial & smooth muscle cells, identifying unique PCs for each cell type specific, light induced state. In contrast, Standard PCA identifies the same PC or state that maximally correlates with light exposure in these two cell types, highlighting the differences in cellular state resolution between each method and the enhanced ability of ResidPCA to identify unique states. These findings underscore the method-dependent nature of state interpretation, as other approaches reveal different patterns (**Supplementary Fig. 2**).

ResidPCA predominantly identifies a distinct light-induced state unique to each cell type, rather than a single light induced state that is conserved across cell types, except for macrophages and endothelial & smooth muscle cell types, which share a single light induced state (**Supplementary Fig. 2**). Moreover, each light-induced, cell type-specific state exhibited distinct Gene Set Enrichment Analysis (GSEA) enrichments compared to those of other cell types. Specifically, we observed that the light-induced state for each cell type had fewer than five overlapping pairwise gene set enrichments with the light-induced states of other cell types suggesting that the light induced states for each cell type differ in functional mechanism (**Supplementary Fig. 2**). Exceptions to this were endothelial & smooth muscle cells and macrophages, which shared the same light-induced state, and interneurons and excitatory neurons, which are very interrelated cell types and often grouped together, which shared greater than seven gene sets (depending on the pathway sets analyzed) (**Supplementary Fig. 2**). These findings indicate that ResidPCA effectively identifies unique light-induced states activated in each cell type and can properly recover distinct cell-type specific states into separate principal components. This observation aligns with the conclusions of Hrvatin et al. that each cell type activates a distinct cell state or set of genes in response to light exposure. While each state was typically associated with distinct gene set enrichments and biological pathways, we observed that many states shared common gene set enrichments related to cellular stress, including pathways such as **upregulated stress responses to photodynamic therapy**, **green/yellow light exposure**, and **chronic stress** (**Supplementary Table 1**). This suggests that while environmental stimuli can trigger distinct state activations in different cell types, these states often converge on core pathways involved in responding to external stressors.

##### Supplementary Note 2, Fig. 2: Cell-type specific stress responses to light exposure: enrichment of conditional stress pathways and gene ontology annotations identified by ResidPCA

Although the light induced state of each cell type is distinct, each of light-induced cell type specific states identified by ResidPCA showed enrichment in conditional stress responses associated with light exposure (Excitatory State: **Chronic Stress in Response to Restraint – Hippocampus** – NES: 1.910, FDR p-value: $p < 1.2e-05$; endothelial/smooth muscle + Macrophage State: **Chronic Stress in Response to Restraint – Hippocampus** – Normalized Enrichment Score (NES): -1.853, FDR p-value: 0.006; Interneurons State: **Upregulated Stress Pathways in Photodynamic Therapy** – NES: 1.836, FDR p-value: 0.002, Mural State: **Green/Yellow Light Exposure** – NES: 2.016, FDR p-value: $p < 1.2e-05$), as well as Gene Ontology enrichments that highlight cell-type specificity (Microglia State: **Embryonic Cortex Microglia** – NES: 1.746, FDR p-value: $p < 1.2e-05$) (**Supplementary Table 1**). Additionally, many GSEA enrichments are consistent with those identified in the cell type-specific states reported in the original publication; 34 out of 488 pathways found in our states overlap with pathways in the Light Hvratin data.

**<u>Supplementary Note 3, Fig. 2:</u> Impact of variable gene selection on state identification accuracy: comparative performance of ResidPCA, Standard PCA, and cNMF**

Selecting the most variable genes is intended to reduce noise from genes with low expression, however the optimal number of variable genes for maximizing state identification accuracy is not well established in the literature, and generally co-opted from cell type identification[84]. In our analysis, using fewer variable genes (3,000) generally improved accuracy, however, ResidPCA was the method least impacted by incorporation of more variable genes (20,000) while cNMF was the most impacted (**Supplementary Fig. 2**). The average change in accuracy when using 20,000 variable genes compared to 3,000 was slightly positive overall (ResidPCA $\Delta R^2 = 0.004$, Standard PCA $\Delta R^2 = 0.018$, cNMF $\Delta R^2 = 0.065$, max $R^2$). The use of 20,000 variable genes showed mixed results, with accuracy sometimes exceeding and sometimes falling short of that achieved with 3,000 variable genes (**Supplementary Fig. 2**). When applying ResidPCA and Standard PCA, certain cell types exhibited decreased accuracy with 20,000 variable genes compared to 3,000 (astrocytes: ResidPCA $R^2 = 0.143/0.114$, Standard PCA $R^2 = 0.118/0.094$; endothelial/smooth muscle: ResidPCA $R^2 = 0.341/0.207$, Standard PCA $R^2 = 0.301/0.152$; mural: ResidPCA $R^2 = 0.273/0.152$, Standard PCA $R^2 = 0.234/0.147$ [3000/20,000 variable genes], joint adj. $R^2$) (**Supplementary Fig. 2**). In summary, ResidPCA exhibited the smallest variability in accuracy relative to the number of variable genes, but no single optimal gene count emerged as universally best, highlighting the need for ongoing research to refine these parameters.

**<u>Supplementary Note 4. Fig 4:</u> Mechanistic analysis of ResidPCA-identified states**

<u>Key ResidPCA cellular states enriched for AD heritability</u>

The pathways enriched for PC15 converged on mechanisms involving the immune system's response to antigenic stimuli–a critical process in the context of AD that intersects the excitatory neuron–oligodendrocyte–microglial axis. One enriched pathway, **antigen processing and presentation of exogenous peptide antigen** (FDR p-val: $p < 1.2e-05$, NES: 2.509), highlights the immune system's role in recognizing and presenting foreign antigens to immune cells, a process vital for initiating an immune response. Similarly, **microglia pathogen phagocytosis pathway** (FDR p-val: $p < 1.2e-05$, NES: 2.507) (**Fig. 4C**) underscores the importance of microglia, the brain's resident immune cells, in phagocytosing pathogens and debris, thus maintaining neuronal health. The **hallmark interferon alpha response** (FDR p-val: 0.004, NES: 1.934) pathway signifies the role of interferons in modulating immune responses, particularly in antiviral defense, which can influence neuroinflammation in AD. The **regulation of inflammatory response to antigenic stimulus** (FDR p-val: 0.004, NES: 1.933) pathway emphasizes the regulation of inflammation upon encountering antigens, a critical factor in the neuroinflammatory processes observed in AD. Lastly, the **regulation of CD4 positive alpha beta T cell differentiation** (FDR p-val: 0.004, NES: 1.932) pathway points to the differentiation of CD4+ T cells, which are pivotal in orchestrating adaptive immune responses. Together, these pathways illustrate a unified mechanism where immune system dysregulation and neuroinflammation play central roles in AD pathogenesis, potentially linking metabolic disruptions and immune responses in the disease's progression. Additionally, interpreting the significant GWAS hits in this state annotation further supports the mechanistic connection to the microglial immune response. The statistically significant AD GWAS gene enriched in this state is RASGEF1C, which has been implicated in microglial activation through immune signaling pathways[53]. Additionally, this state corresponds with a recently identified state that involves both neurons and oligodendrocytes in **beta amyloid production**, particularly evident in early AD. While neurons were already established as producers of amyloid beta, a finding highlighted that oligodendrocytes also produce amyloid beta and share a similar transcriptomic profile with neurons in this context[16]. PC15 in ResidPCA is significantly enriched for a curated gene set associated with **beta amyloid production** (FDR p-val: 0.009, NES: 1.550), spanning the neuron-oligodendrocyte cell type axis. The beta amyloid production by these cell types triggers microglial activation, a subsequent pathogenic event in AD, demonstrating how this state captures the neuron-oligodendrocyte-microglia axis. In summary, this state reflects the production of beta amyloid in both neurons and oligodendrocytes, followed by the activation and response of microglia.

The second most significant ResidPCA-based state for AD heritability, PC27 (Enrichment: 2.001, FDR p-value: 3.589e-08), maps predominantly to metabolism. Upon further inspection of the specific GSEA-related pathways that are enriched, one notable pathway is **drug metabolism cytochrome P450** (FDR p-val: $p < 1.2e-05$, NES: -3.414) responsible for metabolizing drugs and endogenous compounds. Epigenetic markers of genes coding for subproteins of cytochrome P450 have been suggested as AD biomarkers[54]. This protein is a key regulator of cholesterol and xenobiotic metabolism and is implicated in AD[54]. Another highly relevant pathway identified in this ResidPCA state is **steroid hormone biosynthesis** (FDR p-val: $p < 1.2e-05$, NES: -3.207). This pathway converts cholesterol into CYP proteins, which are sub-components of cytochrome P450[55]. Other pathways enriched involving cytochrome p450 as an enzyme catalyzing its metabolic reaction include **retinol metabolism** (FDR p-val: $p < 1.2e-05$, NES: -3.134) and **pentose and glucuronate interconversions** (FDR p-val: $p < 1.2e-05$, NES: -2.929)[56,57]. Together, these enriched pathways suggest a potential mechanism corroborating previous evidence that AD is linked to disruptions in cholesterol and steroid hormone metabolism through cytochrome p450 regulation and steroidogenesis. The statistically significant AD GWAS gene enriched in this state is STX1B, which has not been directly related to metabolism, and is predominantly related to synaptic vesicle exocytosis implicated in neuronal function[85]. The most enriched cell types in this state, although not surpassing the correlation threshold ($R^2$: 0.2), are excitatory neurons and oligodendrocytes (both $R^2$: 0.12) (**Fig. 3 D**). These cell types are strongly affected by hormones and retinol metabolites, which modulate neuronal plasticity and myelin production[58–61]. Additionally, pathways involved in drug metabolism (Cytochrome P450) and carbohydrate metabolism (pentose and glucuronate interconversions) also impact these cells by maintaining their metabolic homeostasis and detoxifying harmful substances, which is essential for their optimal function and survival[56].

A cell type-specific state identified by ResidPCA, uniquely expressed in microglia and enriched for AD heritability, stands in contrast to the previously mentioned states that span multiple cell types. This state is represented by PC44 (Enrichment: 1.814, FDR p-value: 2.509e-05) (**Fig. 4 E**). Our analysis identified three key signaling pathways—**modulators of TCR signaling and T cell activation** (FDR p value: 0.001, NES: 2.159), **CD8 TCR downstream pathway** (FDR p value: 0.002, NES: 2.145), and **differentiating T lymphocyte** (FDR p value: 0.002, NES: 2.162)—that interact within microglia and may contribute to the pathogenesis of AD. The **modulators of T cell receptor (TCR) Signaling and T Cell Activation** pathway, which regulates TCR signaling and subsequent T cell activation, was significantly associated with altered microglial activation states[62]. Dysregulation within this pathway is hypothesized to enhance the inflammatory response in microglia, exacerbating neuroinflammation—a hallmark of AD. Similarly, the **CD8 TCR downstream pathway**, primarily involved in the downstream signaling of CD8+ T cells post-TCR engagement, was also found to be significantly linked to microglial activity[62]. Dysregulated CD8+ T cell signaling, potentially mediated through the secretion of pro-inflammatory cytokines such as IFN-γ and TNF-α, may drive microglia towards a pro-inflammatory phenotype[62]. This shift in microglial activity is associated with synaptic dysfunction and neuronal death, contributing to AD progression. Lastly, the **differentiating T lymphocyte** pathway, which plays a role in T lymphocyte differentiation, further highlights the interplay between peripheral immune signaling and CNS-resident microglia[62]. Collectively, these findings suggest that cross-talk between these immune pathways and microglial activation may be pivotal in the inflammatory processes that underlie Alzheimer's disease.

<u>Key ResidPCA cellular states enriched for AD heritability and anging and/or AD gene set enrichment</u>

The first and only state identified by ResidPCA that was enriched for AD heritability and gene set enrichment for AD and aging was PC6. This state captures a predominantly microglial state ($R^2$: 0.23) (**Fig. 3 D**). This state is significantly enriched for AD heritability (FDR p-value: 2.509e-05), **AD** GSEA (NES: 1.771, FDR p-value: 0.013) and **aging** GSEA (NES: 2.912, FDR p-value: p < 1.2e-05). The GSEA indicates that this microglial state involves microRNAs, both previously associated with AD and novel ones, highlighting the emerging role of microRNAs as biomarkers in AD pathology[86]. Established microRNAs associated with AD and enriched in this state include **MIR221** (NES: -3.062, FDR p < 1.2e-05), **MIR4776_3P** (NES: -3.048, FDR p < 1.2e-05), and **MIR362_3P** (NES: -3.003, FDR p < 1.2e-05), which have been associated with differential expression in AD. Specifically, **MIR221** and **MIR4776_3P** are linked to brain regions, while **MIR362_3P** is associated with cerebrospinal fluid (CSF)[87]. Additionally, this state highlights novel microRNAs not yet linked to AD, including **MIR1322** (NES: -3.301, FDR p < 1.2e-05), **MIR10393_3P** (NES: -3.287, FDR p < 1.2e-05), **MIR767_3P** (NES: -3.280, FDR p < 1.2e-05), **MIR6824_5P** (NES: -3.269, FDR p < 1.2e-05), **MIR4718** (NES: -3.267, FDR p < 1.2e-05), **MIR759** (NES: -3.268, FDR p < 1.2e-05), **MIR6801_5P** (NES: -3.266, FDR p < 1.2e-05), and **MIR6793_3P** (NES: -3.233, FDR p < 1.2e-05).

Further validating the microglial association, GSEA shows enrichment of the pathway **TYROBP causal network in microglia** (NES: 2.896, FDR p-value: p < 1.2e-05). This pathway is critical for microglial activation and inflammatory signaling and activation, involving the TYROBP (DAP12) protein, which interacts with various receptors to modulate microglial responses. The enrichment in this pathway suggests that the identified microglial state is characterized by heightened activation and inflammatory responses, integral to AD pathology[65]. Moreover, this state maps to an inflammatory mechanism, supported by the enrichment of the **TNFA signaling via NFKB** pathway (NES: 2.944, FDR p-value: p < 1.2e-05).

Interpreting the GWAS genes enriched in this state, **HERC1** and Amyloid Precursor Protein (**APP**) reinforce the proposed mechanism. **HERC1**, involved in the ubiquitin-proteasome system, plays a role in protein degradation and immune responses, potentially influencing inflammatory signaling pathways[88]. **APP**, a key player in Alzheimer's disease, is known to activate microglia and contribute to neuroinflammation through its cleavage products, such as amyloid-beta[89,90]. Together, these genes suggest a pathway where dysregulation in protein degradation and amyloid processing triggers microglial activation and inflammatory responses, contributing to AD pathology. This mechanism underscores the critical role of microglia and inflammation in the progression of Alzheimer's disease, mediated through both established and novel microRNAs.

Shifting focus from states enriched in both AD heritability and gene sets related to **AD** and **aging**, we now examine states with AD heritability enrichment combined with only **aging**-related gene set enrichment. These states may be less specific to the disease but more closely associated with aging. PC3 is a state identified by ResidPCA that contains enrichment for AD heritability and GSEA enrichment for aging (NES: -3.365, FDR p-value: p < 1.2e-05 (**Fig. 4 E**). The orientation of this state can be determined by observing that AD heritability enrichment is significant in the genes on the positive axis of the embeddings (FDR p-value: 0.024). This indicates that the diseased state is associated with cells positioned along the positive end of the embedding continuum. The mechanistic interpretation of the cell state involving astrocytes ($R^2$: 0.38), microglia ($R^2$: 0.25), and oligodendrocyte ($R^2$: 0.35) (**Fig. 3 D**) reveals significant disruptions in glial cell function, particularly impacting oligodendrocytes, and connects these disruptions to AD pathology. The NES for the pathway **oligodendrocyte markers** (NES: -4.151, FDR p-value: p < 1.2e-05) indicates a substantial reduction in markers associated with mature oligodendrocytes. This suggests an impairment in the differentiation and function of these cells, which are crucial for maintaining myelin integrity in the CNS. This impairment is further corroborated by a downregulation in the **cholesterol biosynthesis** pathway (NES: -2.928, FDR p-value: p < 1.2e-05), reflecting decreased cholesterol biosynthesis. Cholesterol is a key component of the myelin sheath, and its reduced production can compromise myelin formation and function.

Additionally, the negative NES in **axon ensheathment in CNS** (NES: -2.811, FDR p-value: p < 1.2e-05) highlights a diminished ability of oligodendrocytes to properly ensheath axons, leading to impaired neural signal transmission. This finding is particularly relevant to AD, where axonal damage and myelin loss are prominent features. The disruption of myelin formation and maintenance can contribute to the progressive neurodegeneration observed in AD.

The negative NES in **protein refolding** (NES: -2.589, FDR p-value: p < 1.2e-05) suggests that cellular stress and protein misfolding are occurring within glial cells, potentially exacerbating the pathological processes in AD. Protein misfolding and aggregation are hallmarks of AD, particularly with respect to amyloid-beta and tau proteins. This protein stress within glial cells could further impair their function and contribute to disease progression.

The decreased activity in **regulation of glial cell differentiation** (NES: -2.272, FDR p-value: 0.004) indicates impaired regulation of glial cell differentiation, which affects the maturation of oligodendrocyte precursor cells (OPCs) into fully functional oligodendrocytes. Given that oligodendrocytes are crucial for myelination and neural support, their dysfunction can exacerbate the neurodegenerative effects seen in Alzheimer’s disease.

The association with **APP (Amyloid Beta Precursor Protein)**, a key gene linked to Alzheimer’s disease, further underscores the relevance of these glial disruptions to AD pathology. **APP** is involved in the production of amyloid-beta, which forms plaques in Alzheimer’s disease, and activates glial cells, which forms plaques in AD and triggers the activation of glial cells responsible for clearing these misfolded protein aggregates[91]. Ineffective clearance of these aggregates results in neuroinflammation and neuronal damage while myelin dysfunction implicated in reduced axon formation has been associated with amyloid-beta deposition leading to AD pathology[92].

Finally, another state enriched in both AD heritability (FDR p-value: 8.472e-03) and the **aging** gene set (NES: 1.680, FDR p-value: 0.032), but identified in IterPCA for MG rather than ResidPCA, is PC4 of IterPCA in MG. The identified cell state involving the AD GWAS genes **RHOH**, **C14orf80**, and **LIME1** is found exclusively in microglia, highlighting a significant mechanistic role for these cells in AD. **RHOH** and **LIME1** are integral components of T-cell receptor (TCR) signaling pathways, which is an enriched gene set (**TCR signaling pathway** – NES: 2.628, FDR p-value: p < 1.2e-05). This pathway is crucial for microglial cell activation and signaling[62,93]. In microglia, the brain's resident immune cells, these signaling pathways can modulate the cells' responses to neuronal damage and amyloid-beta plaques, which are characteristic of AD[94].

Microglial activation via TCR signaling pathways can lead to the production and release of pro-inflammatory cytokines, exacerbating neuroinflammation—a key feature of AD[62]. Chronic activation of microglia can result in a sustained inflammatory state, contributing to neuronal injury and synaptic loss[94]. The presence of **C14orf80** within this microglial state is unclear.

Thus, the mechanistic interpretation of this state underscores the importance of microglial activation and adhesion in the pathogenesis of AD. The persistent activation of microglia through T-cell receptor signaling pathways, highlights how immune dysregulation within these cells can drive neuroinflammation and contribute to the progression of AD.

**Supplementary Note 5. Fig 5: Efficiency comparison of ResidPCA Toolkit and Seurat for ResidPCA implementation**

We evaluated the efficiency of performing ResidPCA using the ResidPCA Toolkit compared to an implementation of the same method using Seurat. Notably, ResidPCA implemented with the ResidPCA Toolkit was more efficient, requiring less CPU time (mean CPU time: 0.15 hours, SE: 0.037) and memory (mean max RSS: 79.28 GB, SE: 12.77) than the implementation with Seurat (CPU time: 6.02 hours, SE: 0.43; max RSS: 111.60 GB, SE: 4.40) (**Fig. 5 C)**. For instance, applying ResidPCA to a count matrix with 100,000 cells and 20,000 genes, the ResidPCA Toolkit completed the task in 0.32 hours (SE: 0.03), whereas Seurat required 15.12 hours (SE: 0.54)—a 47-fold increase in time (**Fig. 5 C)**. Additionally, when processing a dataset with 25,000 cells and 10,000 genes, performing ResidPCA with the ResidPCA Toolkit utilized one-third as much maximum memory than implementing ResidPCA with Seurat (11.47 GB, SE: 3.01 vs. 35.96 GB, SE: 0.04). Both CPU time and memory usage scaled linearly with the number of genes in the count matrix while keeping the number of cells constant (**Fig. 5 A, B, D**).
